# A genomic exploration of the possible de-extirpation of the Zanzibar leopard

**DOI:** 10.1101/2023.07.25.550323

**Authors:** Xin Sun, Emily Louisa Cavill, Ashot Margaryan, Jianqing Lin, Søren Thingaard, Tamrini A. Said, Shyam Gopalakrishnan, M. Thomas P. Gilbert

## Abstract

The recently extirpated Zanzibar leopard was the only known African leopard (*Panthera pardus spp.*) population restricted exclusively to a major island habitat. Although its demise was driven through habitat change and conflict with humans, given its role as a keystone species for the Zanzibar Archipelago, its potential reintroduction might offer a means for helping preserve the natural biodiversity of its former habitat. Whether this is feasible, however, would be contingent on both whether closely related source populations can be identified on mainland Africa, and whether the Zanzibar form exhibited any special adaptations that might need to be considered when choosing such a source. In light of these questions, we genomically profiled two of the six known historic specimens, to explore whether they represent a realistic candidate for de-extirpation through reintroduction. Our analyses indicate that despite its geographic separation, the Zanzibar leopard shared a close genetic relationship with mainland East African individuals. Furthermore, although its uniqueness as an island population was emphasised by genomic signatures of high inbreeding and increased mutation load, the latter similar to the level of the critically endangered Amur leopard (*P. p. orientalis*), we find no evidence of positive selection unique to Zanzibar. We therefore conclude that should attempts to restore leopards to Zanzibar be planned, then mainland East African leopards would provide a suitable gene pool, whether using genetic engineering or conventional rewilding approaches.

## Introduction

Biodiversity losses resulting in both the extirpation of unique populations, and extinction of whole species, have intensified in past decades despite the increase of conservation efforts (Butchart et al., 2010; Pereira et al., 2010; Seddon et al., 2014a). To restore lost diversity and ecosystem functions, the validity of a range of different conservation approaches are being debated, and even implemented, into various ecosystems through different approaches (Fernandez et al., 2017; Perino et al., 2019; Seddon et al., 2014a; Svenning et al., 2016). For example, on the one hand, given recent technological advances in genome editing and the retrieval of genetic information from Pleistocene megafauna (Dabney et al., 2013; Gansauge and Meyer, 2013), the idea of de-extinction as a creator of proxy for lost biodiversity in order to restore ecological function is, while heavily debated, gaining traction (IUCN/SSC, 2016; Lin et al., 2022; Novak, 2018; Richmond et al., 2016; Seddon et al., 2014b; Shapiro, 2017). On the other hand, technologically simpler methods are also being explored, including back-breeding to restore lost phenotypes or even lost species (“The Quagga Project,” n.d., “The Tauros programme,” n.d.), as well as reintroduction of extirpated populations from closely related gene pools (Lorimer et al., 2015).

To date, relatively few successful examples have been delivered using these approaches. Of these, reintroduction approaches have delivered the most success in recent years, including species spanning diverse spatial, temporal and genetic backgrounds (Germano and Bishop, 2009; Griffiths et al., 2013; Keedwell et al., 2002). For example, the rewilding of the wolf population in the Yellowstone National Park has played the role of a keystone species (Lorimer et al., 2015; Ripple and Beschta, 2012) and reshaped the local ecosystem, while the replacement of an extinct, with a related extant, tortoise species has been used to fill an otherwise vacant niche in the Galapagos Islands (Hunter et al., 2013; Lorimer et al., 2015). In a different context, through back-breeding, at least morphologically (if not genomically) similar quagga-like plain zebras have been regenerated through selective crosses between closely related zebra subspecies (Larison et al., 2021; “The Quagga Project,” n.d.). And lastly, although still in its early stages, genome editing technologies such as CRISPR, have boosted a growing interest in ambitious projects that aim to resurrect now lost megafauna, such as the woolly mammoth (“ColossalTM/ A New Dawn of Genetics,” n.d.), and the recently extinct Tasmanian tiger (“ColossalTM/ A New Dawn of Genetics,” n.d.).

Ultimately, for any of these approaches to gain favour, test cases are needed to explore whether the reality matches the expectations, and success stories are needed to showcase the methods. In this regard, it could be argued that ideal candidates are those that have perceived conservation benefits, while also being technically feasible, and likely to generate sufficient support from the public (IUCN/SSC, 2016; Seddon et al., 2014b). Furthermore, in addition to the technical limitations associated with resurrection attempts (for example relating to integrity of the candidate genome), analysis of risks for possible impacts on both the ecology and human society of the region that would receive the reintroduced or resurrected species needs to be performed (Iacona et al., 2017; Seddon et al., 2014b).

The Zanzibar leopard may offer an interesting candidate for de-extirpation through reintroduction, for several reasons. As a morphologically distinct, and now extirpated, population of a charismatic big cat species, the leopard (*Panthera pardus*), it holds not only public appeal but also belongs to a species that exhibits extraordinary adaptation to versatile habitats, something critical should animals require introduction to novel habitats. Furthermore, surviving African leopards maintain relatively high genomic diversity, coupled to low levels of differentiation, in comparison to other big cats (Paijmans et al., 2021; Pečnerová et al., 2021). Unfortunately however, while once endemic to Unguja Island, upon which it was the apex predator, ultimately the Zanzibar leopard failed to survive the challenge of human colonisation of the Zanzibar archipelago (Walsh and Goldman, 2007), likely being fully extirpated by the end of the 20th century (Goldman and Walsh, 2002) due to extensive hunting. With regards to phenotype, in comparison to its continental relative, this apex predator is thought to have had a smaller body size and distinct skin pattern (Pakenham, 1984). With no captive individuals known, six preserved specimens collected in the early 20th remain the only source from which to study this population (Walsh and Goldman, 2008).

## Results and Discussion

### Whole-genome sequencing of the Zanzibar leopard

Three of the six known Zanzibar leopard specimens (Walsh and Goldman, 2008) were sampled for this study, including one sample of hairs from the specimen held on display at the Zanzibar Museum (Z 1209) that dates to the first half of the 20th century (Goldman and Walsh, 2002), and cuttings of skin and dried tissue from both specimens held at the Harvard Museum of Comparative Zoology (MCZ36709, 1937 and MCZ40953, 1939). DNA was extracted and sequenced from all three samples. The sample from the Zanzibar museum specimen failed to deliver any usable sequencing libraries, thus was discarded from further analysis. In contrast, both samples from Harvard yielded rich libraries containing relatively high endogenous DNA levels (70.7% for MCZ36709 and 38.8% for MCZ40953). Given both the similarity in age of the two specimens, and the results of an initial analysis revealing they carried nearly identical mitogenome sequences (Fig. S1), we elected to focus subsequent sequencing on the sample with best quality DNA (MCZ36709), resulting in final average genomic depth of coverage of 38.4 ×. Next, to clarify the genetic ancestry of the Zanzibar leopard, a genomic dataset representing the genetic background of modern and historic leopards (average sequencing depth of 9×) was assembled from published studies (Paijmans et al., 2021; Pečnerová et al., 2021). Subsequent analyses were restricted to transversions only, to account for possible DNA damage derived errors in the resulting dataset, thus yielding a final matrix containing 2,156,661 transversions, representing 80 African and Asian leopard samples (Fig. 1A, Table S1).

**Figure 1.**
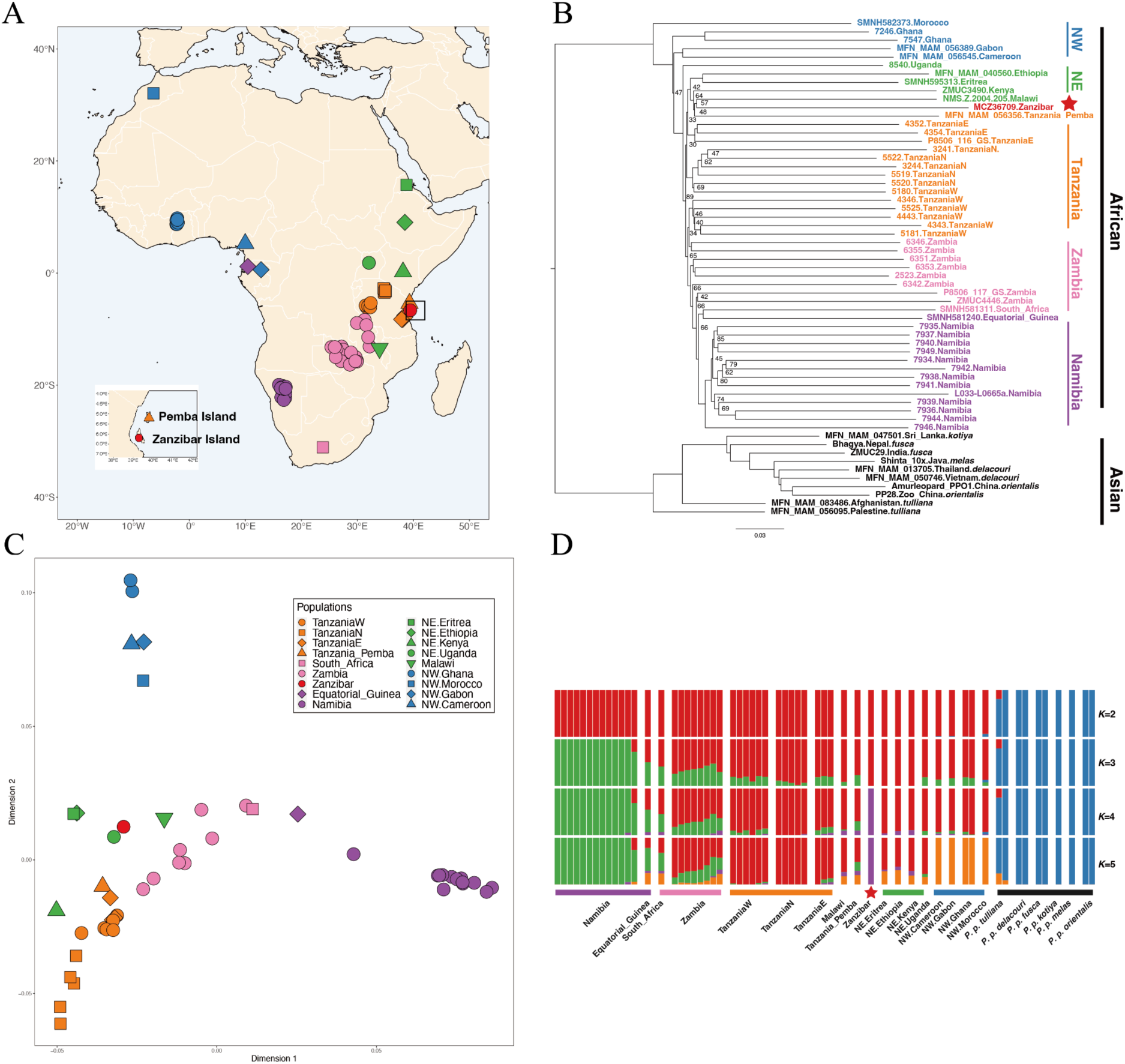
Population structure of African leopards. (A) Geographic origin of the Zanzibar leopard and other African leopard samples (Table S1). (B) Whole-genome phylogeny inferred using Neighbour-Joining methods. Bootstrap support values are shown for nodes with values lower than 90. (C) Multidimensional scaling plot portraying the genomic diversity of African leopards. Samples are labelled with their geographical origin. Icons in A-C were coloured based on their geographical proximity. (D) ngsAdmix plot assuming two to five different ancestries of Asian and African leopards. African leopard (*P. p. pardus*); Amur leopard (*P. p. orientalis*); Javan leopard (*P. p. melas*); Sri Lankan leopard (*P. p. kotiya*); Indian leopard (*P. p. fusca*); Indochinese leopard (*P. p. delacouri*); Persian leopard (*P. p. tulliana*).

### The Zanzibar leopard is a unique island population of the African leopards

To explore the genetic affinity between the Zanzibar and other leopards, we first looked at the population structure of our dataset. The results (Fig. 1 B-D and Fig. S2) both recapitulate previous observations of continent level separation in leopards (Paijmans et al., 2021; Pečnerová et al., 2021), and reveal that the Zanzibar leopard falls within the diversity of African leopards. Subsequently, we restricted multidimensional scaling (MDS) analysis to African leopards, and observed that the placement of the Zanzibar leopard is distinct from the major Eastern and Southern population clusters of Zambia, Namibia, and Tanzania (Fig. 1B). Additional analysis in a whole-genome phylogeny revealed that despite this separation, the Zanzibar leopard forms a clade with samples from Northeast Africa (refer to as NE population), including Kenya, Eritrea, and Ethiopia (Fig. 1C), but not, perhaps surprisingly, with mainland Tanzania, where its nearest geographical relatives are found. We highlight, however, that although clusters are formed within African leopards in the phylogeny, the node support for each clade was not high, possibly due to gene flow between these populations and/or incomplete lineage sorting due to a recent divergence time (Pečnerová et al., 2021). However, while estimating the individual admixture proportions for each leopard sample at *K*=5, the Zanzibar leopard formed a distinct cluster (Fig. 1D).

### Low levels of divergence between the mainland and island population

As the Zanzibar leopard is most likely extinct in the wild, and because no reliable records exist of captive individuals (Goldman and Walsh, 2002), the gene pool of mainland African leopards would be the most suitable source for any future re-introduction attempt. We explored this by asking which mainland African population has the closest relationship with the Zanzibar leopard. Specifically, *f*3-statistics were used to compare the shared ancestry between the Zanzibar and other leopards (Fig. 2A). Surprisingly, instead of finding a closest relationship with the geographically most proximate East Tanzanian population, most ancestry was shared with samples from the more North-Eastern population (Eritrea and Kenya), along with a sample from much more to the South, in Malawi. Also notable was the observation that a previously sequenced sample labelled as originating from Pemba Island within the Zanzibar Archipelago, is more closely related to the mainland Tanzanian samples than our Zanzibar specimen. Given that a previous study has questioned the reliability of the geographic origin assigned to the Pemba Island sample (Paijmans et al., 2021), we cannot rule out that it may truly derive from the African continent. Furthermore, we cannot exclude that similar concerns about geographic provenance errors may explain the unexpectedly close relationship of the Zanzibar specimen with that from Malawi.

**Figure 2.**
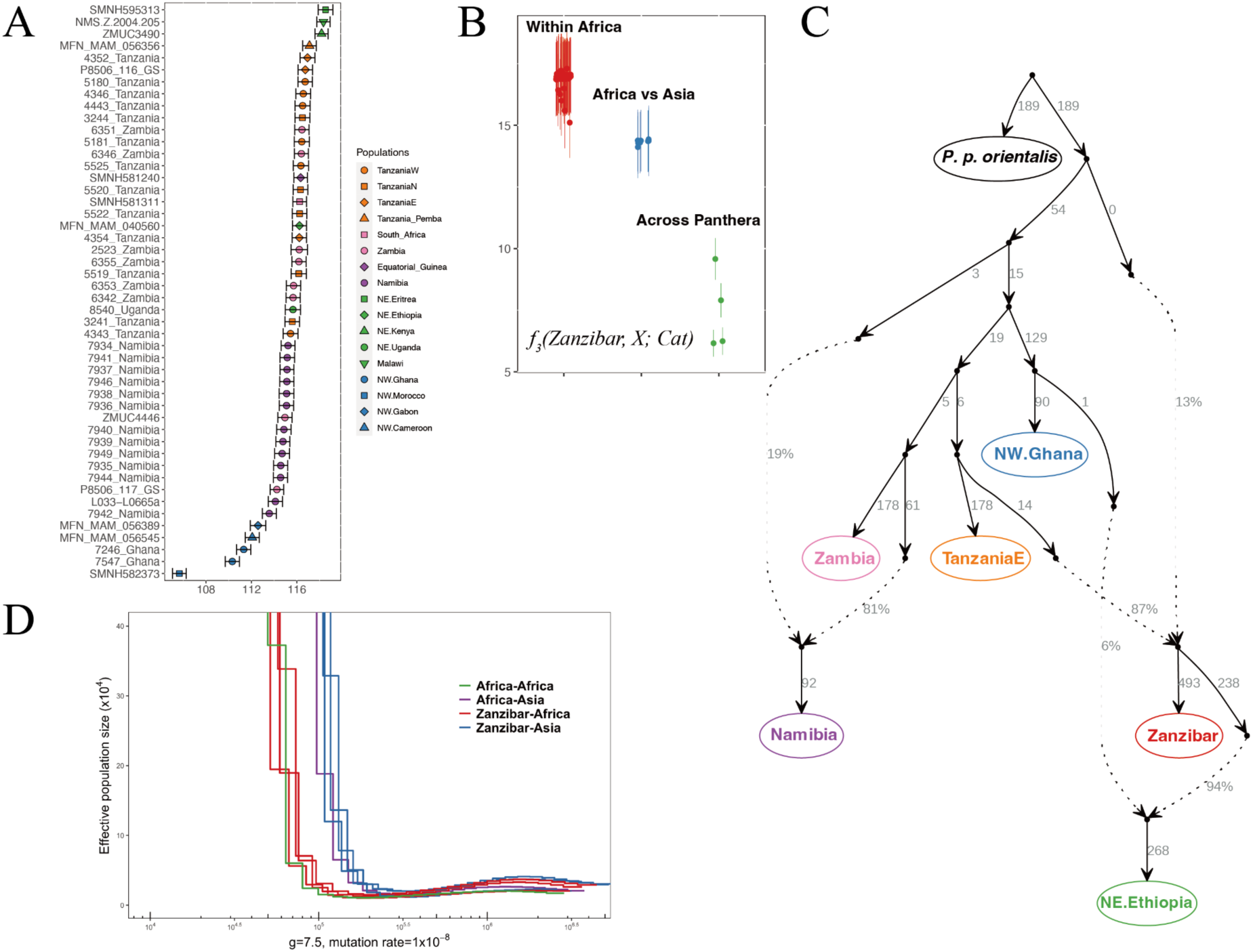
Genetic affinity between the Zanzibar and mainland African leopards. (A) Outgroup *f*3-statistics comparing shared ancestry between the Zanzibar leopard and mainland African leopards. Icons are coloured and shaped according to their geographical and genetic clusters. (B) *f*3-statistics comparing shared ancestry between the Zanzibar leopard and all *Panthera* samples included in this study. (C) Admixture Graph modelling the ancestry of African leopards. The Zanzibar leopard was modelled to have ancestry (13%) from an early diverged African leopard lineage. (D) Divergence dating by hPSMC showing a similar level of divergence between the Zanzibar leopards and mainland African leopards and within mainland African leopards. A pseudo-diploid X chromosome of two male leopards excluding transversion sites was used assuming a generation time of leopards as 7.5 years (Pečnerová et al., 2021) and the mutation rate as 1 x 10^-8^ per nucleotide per year (Figueiró et al., 2017).

In general, the level of the shared ancestry between the Zanzibar and other samples is substantially higher than both that seen when comparing leopards from the two continents that they inhabit, or with other *Panthera* species, which supports a general pattern of low-level divergence within the African continent (Fig. 2B). We subsequently validated the divergence level by estimating relative divergence times using hPSMC. To do this, hybrid leopard X chromosomes were generated and compared between pairs of male individuals, including (i) the Zanzibar leopard and one other African leopard, (ii) the Zanzibar leopard and one other Asian leopard, (iii) two randomly selected African leopards to represent within Africa divergence level, and (iv) one African leopard and one Asian leopard to represent inter-continental divergence. Our analyses reveal that the estimated divergence time between the Zanzibar leopard and other African leopards is similar to the divergence time within the mainland African leopards, possibly due to the isolation caused by glacial period cycles in the Late-Pleistocene (Prendergast et al., 2016). Although the Zanzibar leopard and the mainland African leopard shared similar divergence times, we identified an extremely low number (less than 0.005) of identical-by-descent (IBD) segments shared between the Zanzibar leopard and its mainland relatives, something that would be consistent with a lack of recent gene flow between the two groups (Fig. S2).

Next, to get a comprehensive understanding of its ancestry, we fitted the Zanzibar leopard and other African leopards into admixture graphs based on *f4*-statistics. Using a heuristics search, the best-fitted graphs modelled an admixed ancestry of the Zanzibar leopard that includes around 10% ancestry from an early diverged branch, with the rest related to the East Tanzanian leopards. The close affinity with the NE population was also modelled as the Ethiopian sample having a large proportion of its ancestry from the sister clade of the Zanzibar leopard.

### Low genetic fitness and lack of unique genomic adaptation signals to the island

Considering its close genetic relationship with the NE population, we hypothesise that the apparent morphological uniqueness of the Zanzibar leopard resulted from its existence as an island population. In general, island mammals populations tend to exhibit less diversity than mainland populations (Frankham, 1997), thus we investigated whether this was the case by comparing the level of genetic fitness-related parameters in the Zanzibar leopard versus other leopards (Fig. 3). In comparison to other African leopards, the Zanzibar leopard showed a much lower level of individual genome-wide heterozygosity, at about one-third of the level of its mainland neighbours. A higher inbreeding coefficient estimated from long runs of homozygosity was also estimated for the Zanzibar leopard, with over 60% of its genome lacking heterozygosity, which in turn gave it an increased mutation load (Fig. 3). Overall, therefore, the genetic fitness level of the Zanzibar leopard was comparable to the currently critically endangered Amur leopards (*P. p. orientalis*) (Paijmans et al., 2021; Uphyrkina et al., 2002), although it is striking that this was already the case in the Zanzibar leopard at time of collection in the 1930s, something that might have been caused by long term conflicts with humans within the geographically restricted area of Unguja island (Goldman and Walsh, 2002).

**Figure 3.**
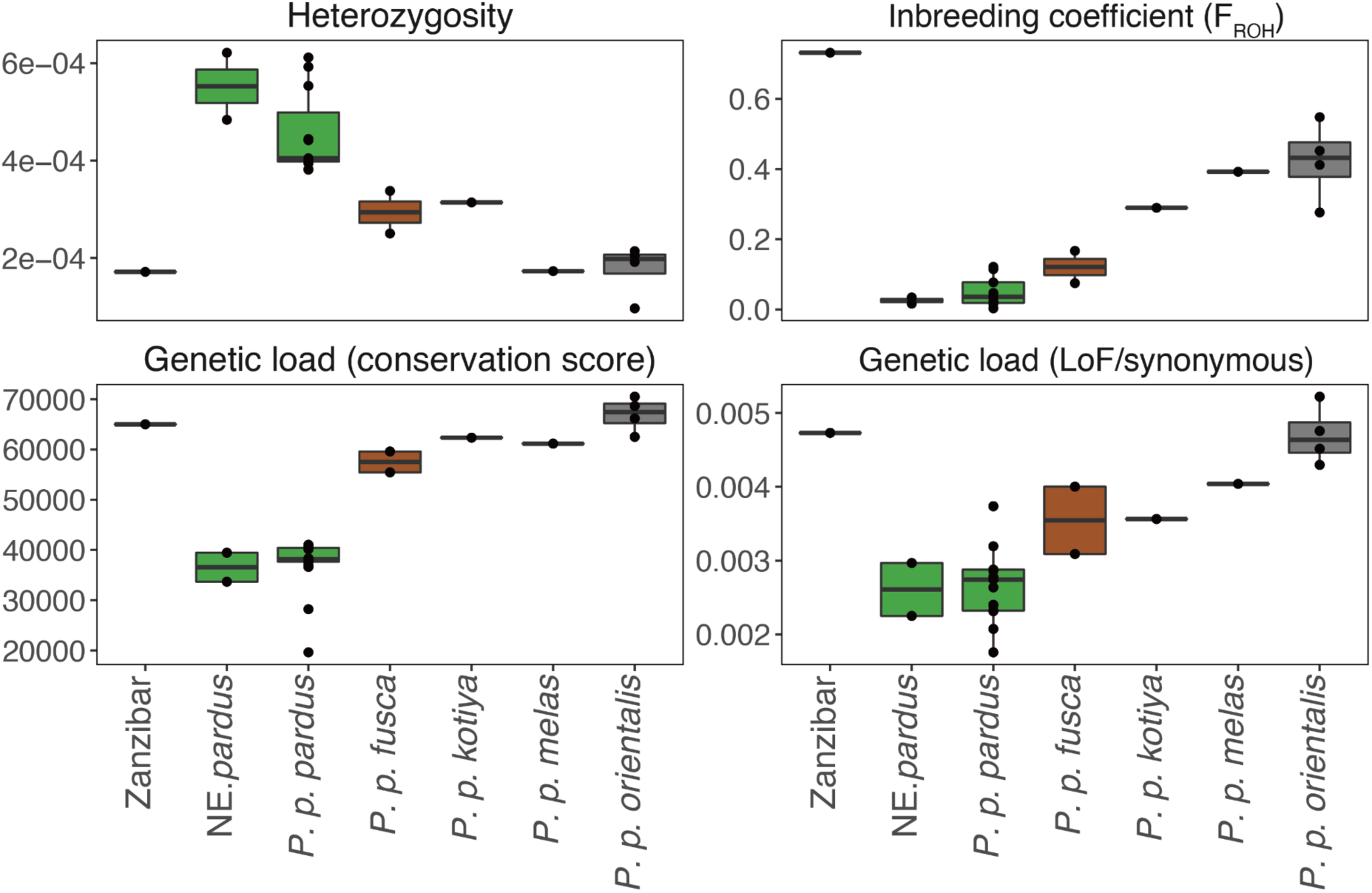
Genome-wide heterozygosity, inbreeding coefficient (F_ROH_), and genetic load for different leopards. Samples are grouped by different leopard subspecies, with the Zanzibar leopard and the northeast African population shown separately. Heterozygosity was calculated from the SFS estimated by ANGSD for each sample. Inbreeding coefficients (F_ROH_) were calculated as the proportion of the genome covered by long runs of homozygosity (ROH ≥ 1Mb). Genetic load was calculated as the sum of high PhyloP scores (top 5 percent across the whole-genome) of homozygous derived transversion sites, and the ratio of loss of function(LoF)/synonymous homozygous derived transversion sites separately. African leopard (*P. p. pardus*); Amur leopard (*P. p. orientalis*); Javan leopard (*P. p. melas*); Sri Lankan leopard (*P. p. kotiya*); Indian leopard (*P. p. fusca*); Indochinese leopard (*P. p. delacouri*).

To explore the possible effects of an increased mutation load, we annotated the Gene Ontology and KEGG pathways of all the genes with an increased level of homozygous derived loss of function and non-synonymous alleles specific to the Zanzibar leopard. Only one pathway, “Bile secretion” (FDR adjusted p-value 0.037), was significantly enriched in this list (Table S2). Although we speculate that this could be related to the difference in food resources between the Zanzibar archipelago and mainland Africa, we caution that both only one individual was included in this analysis, and that leopards are known to have a broad dietary niche (Athreya et al., 2016; Havmøller et al., 2021). We additionally explored for a possible genetic basis for the Zanzibar leopard’s reduced body size and unique coat patterns (Table S3), and identified that a number of genes related to body size (Chase et al., 2002; Hayward et al., 2016; Hoopes et al., 2012; Jones et al., 2008; Parker et al., 2009; Plassais et al., 2022, 2019; Rimbault et al., 2013; Sutter et al., 2007; Vaysse et al., 2011), such as *SMOC2* and *LCORL,* and coat colouration patterns (*MC1R*) (Eizirik et al., 2010, 2003; Kaelin et al., 2021; Peterschmitt et al., 2009) exhibit homozygous non-synonymous alleles in the Zanzibar sample. Interestingly, while these may explain the phenotype, we note that none of them are unique to the Zanzibar leopard population, but rather vary in allele frequency in the mainland African leopards, which could be relevant to consider if attempts were to be made to restore the lost phenotype.

### Re-introduction as a means to de-extirpate the Zanzibar leopard

In light of our results, we propose that there are several grounds that render the Zanzibar leopard a suitable candidate for possible de-extirpation on Unjuga Island. Firstly, although the Zanzibar leopard was unique with regards to its island habitat and morphology, relatively closely related populations remain on the mainland that notably contain genetic variants that may be relevant for its unique morphology. Secondly, as the former apex predator of Unjuga island (Walsh and Goldman, 2007), its reintroduction could have a significant impact on the island’s ecology. Lastly, from a practical point of view, leopards are by far the most successful of the big cats in terms of the types of general survival and adaptability, thus as long as sufficient protection measures are taken to prevent their hunting, it seems likely that any re-introduced leopards would be able to adapt to their new habitat.

In light of these points, if de-extirpation was to be attempted by the relevant local authorities, how might it be achieved? On the one hand, a relatively straightforward solution could be the introduction of mainland African leopards from NE Africa onto the islands. On the other hand, more sophisticated solutions could also be considered, aimed at generating a leopard more closely resembling the original form, using the genomic information generated in our study. In this context, currently three principal methods exist: back-breeding, cloning, and genetic engineering (Richmond et al., 2016; Shapiro, 2015). Back-breeding approaches involve, at the simplest level, selective breeding to reacquire the lost target’s phenotypic traits, as has been pioneered in the context of the quagga (*Equus quagga*) (“The Quagga Project,” n.d.) and the aurochs (*Bos primigenius*) (Stokstad, 2015; “The Tauros programme,” n.d.). Therefore attempts could be made to breed for the reduced size and unique coat morphology using carefully chosen morphological variants found on mainland Africa. At a more advanced level, should extant relatives retain partial genomic fragments derived from the lost form (as has been for example reported in some modern cattle breeds for the aurochs (Park et al., 2015)), then a more nuanced form of back-breeding could be attempted. Specifically, living relatives chosen based on their genomes containing different genomic tracts derived from the lost form could be crossed, ultimately deriving offspring that contain incrementally more of the lost genome (Sinding and Gilbert, 2016). Unfortunately however, given we found no evidence of significant IBD tracts shared between the Zanzibar and mainland African leopards, this does not seem a feasible solution. The second general approach, cloning, is unfortunately not an option, given that no known living Zanzibar leopards exist from which to source the viable cells needed. The final option, genetic engineering, will become increasingly interesting as the technical challenges that it currently faces are resolved (“Colossal^TM^/ A New Dawn of Genetics,” n.d.). Such an approach could both allow direct editing of genomes derived from extant continental African leopard to introduce any genetic variants identified in the Zanzibar leopard genome that were putatively relevant for its morphology and inhabiting its niche, while also ensuring that sufficient genetic diversity exists within the recreated population to enable them to maintain a genomically healthy population.

### Limitations of the study

In summary, deep sequencing of a Zanzibar leopard genome has enabled us to resolve its genetic ancestry, and in doing so provide suggestions for possible routes to its de-extirpation, should the local community on Zanzibar one day wish to explore this option. Naturally, an obvious caveat is that as so few specimens remain for study, our analyses may not reveal a fully representative insight. However, we feel this is unlikely considering the levels of inbreeding observed.

## Author contributions

M.T.P.G., S.T., T.A.S. and X.S. conceived the study. E.L.C. performed the ancient DNA lab work. X.S., A.M., E.L.C. and J.Q.L. performed bioinformatic analysis with input from S.G. and M.T.P.G. S.T. and T.A.S. provided samples from the Zanzibar Museum. X.S. wrote the manuscript with input from all authors.

## Acknowledgements

We thank Salim Kitwana Sururu, Maneno I. Khamis, The Department of Museum and Antiquities, Zanzibar, the Harvard Museum of Comparative Zoology Mammalogy Collection and MCZ-CRYO for providing specimens of the Zanzibar leopard. We also thank Rasmus Heller and Ida Moltke for early access to their African leopard genomic dataset, and Xiao Xu from Peking University for his input on candidate genes of phenotypic traits in the Zanzibar leopards. We thank Sarah S. T. Mak for her help with lab work. This project was funded through ERC Consolidator Award 681396 ‘Extinction Genomics’ awarded to MTPG.

## STAR methods

### Sample collection, DNA extraction, and sequencing

Samples were collected from three of the known Zanzibar leopards: one from the Zanzibar Museum specimen provided under permit IMMK/MKM/68/VOL.IV/36 (sample ID Z 1209, dating to the first half of the 20th century (Walsh and Goldman, 2008)), and one each from each of the specimens held at the Harvard Museum of Comparative Zoology under permit 2020-1-Cryo (sample IDs MCZ36709, 1937 and MCZ40953, 1939).

DNA was extracted in an aDNA-dedicated PCR-free laboratory, using aDNA methods following (Dabney et al., 2013; Mak et al., 2017; Meyer and Kircher, 2010; Ramos-Madrigal et al., 2021). We were unable to construct useable libraries from Z 1209, thus only MCZ36709 and MCZ40953 were successfully converted into BGISeq-compatible genomic libraries using a tailored version of the Santa Cruz Reaction (SCR) (Kapp et al., 2021) single-strand library construction protocol. Libraries were purified using Qiagen MinElute reaction clean-up columns. Four 3uL replicates from each library template were amplified following an in-house uracil-tolerant protocol containing 2.5U PFU Turbo CX Polymerase, 1x PFU Turbo buffer, 0.4 mg ml^-1^ bovine serum albumin (BSA), 0.25 μM mixed dNTPs, 0.1 μM BGI forward primer, 0.1μM BGI reverse index-primer per 50uL reaction. Initial denaturation of libraries was carried out at 95°C for 2 minutes, followed by 30 cycles of denaturation at 95°C for 30 seconds, annealing at 60°C for 30 seconds, and extension at 72°C for 110 seconds, and the final extension held at 72°C for 10 minutes, and the PCR product was then purified with MagBio HiPrep PCR clean-up beads. Amplified libraries were initially sequenced on the DNBSEQ-G400 platform using BGI Copenhagen’s commercial service. In order to generate a high coverage genome for this species, we selected the sample with the highest endogenous DNA content for deeper sequencing (MCZ36709). Subsequently, five additional Illumina libraries were built following the SCR protocol. Three 5uL amplification replicates (following the PFU protocol as above, using Illumina indexes, and 16 thermal cycles per replicate) were made from each of the five new library templates, and were pooled and sequenced across one Novaseq S4 lane at the GeoGenetics Sequencing Core, Copenhagen.

### Data mapping and dataset preparation

We used the *Paleomix* v1.2.13.2 (Schubert et al., 2014) pipeline to map the Zanzibar leopard sequencing data, and data from 79 previously published leopards (Paijmans et al., 2021; Pečnerová et al., 2021), against the domestic cat (*Felis catus*) reference genome (felCat 9.0, RefSeq assembly accession: GCF_000181335.3).

Genotype likelihoods were estimated with ANGSD (Korneliussen et al., 2014) using the GATK model (-GL 2), using reads with mapping quality above 20 (-minMapQ 20) and bases with sequencing quality above 20 (-minQ 20). Sites with a minor allele frequency of less than 0.05, or with more than 50% of the samples having missing genotype information were removed. In order to minimise the effect of aDNA-induced errors in the data generated on the historical samples, we removed from subsequent analyses all transversions. This resulted in a genotype likelihood panel containing 6,642,796 sites, that we refer to henceforth as the genotype likelihood panel.

We also generated a pseudo-haploid panel using ANGSD, by randomly sampling one read (-dofasta 1). Only samples that passed error rate filtering, with relatedness level above 1st degree and a minimum sequencing depth above 5× were included to determine the variable sites. The following command line was used to generate the pseudo-haploid fasta sequence for each sample: -dofasta 1 -doCounts 1 -minQ 20 -minmapq 20 -setminDepthInd 1 -remove_bads 1 -uniqueOnly 1. The variable site panel for all samples was then converted to Plink files for further filtering. We used Plink v2.0 (Chang et al., 2015) to filter the panel with a minimum allele frequency of 0.05 (--maf 0.05), and removed variants in linkage disequilibrium (LD) within a 10kb window, using the correlation threshold of 0.5 (--indep-pairwise 10kb 2 0.5). Transversion sites and sites with repeat sequence annotations were removed. This resulted in a pseudo-haploid panel with 2,156,661 sites, representing 57 samples, that we henceforth refer to as the haploid panel.

We called genotypes for samples that had a sequencing depth above 8× and that passed the quality control. Samtools v1.4 (Li et al., 2009) was used for genotype calling with the following parameters *-Q 20 -q 20 -B --ff 260*. Only biallelic SNPs were kept. To reduce genotype calling error, sites with sequencing depths of lower than 5×, and heterozygous genotypes with sequencing depths for each allele of less than 3×, were masked as missing.

### Population structure with PCA, MDS and NGSadmix

Principal component analysis (PCA) was performed using PCAngsd (Meisner and Albrechtsen, 2018) on the genotype likelihood panel. We conducted two separate PCA analyses, one with all 59 samples that passed the quality control and relatedness filter, and another with only the 49 African leopards that had also passed the same filtering criteria (Table S1).

To generate the multidimensional scaling plot (MDS), we first calculated a pairwise IBS distance matrix using Plink v1.9 (Chang et al., 2015) (--distance square ibs flat-missing), and then used the *cmdscale* function in R to perform dimensional scaling. MDS was conducted with the haploid panel and with the same two separate analyses as PCA.

To infer population structure and admixture for each sample, we used NGSadmix (Skotte et al., 2013) with the genotype likelihood panel of 59 samples as input (Table S1). We ran the analysis assuming 2-6 ancestral populations. Each run was performed independently with 50 replicates and checked manually to assure the convergence of the result.

### Genetic affinity with outgroup f3-statistics and IBD inference

To obtain the shared ancestry between the Zanzibar and other African leopards, we calculated outgroup *f3*-statistics. The Amur leopards were used as the outgroup, and the shared ancestry was measured since the separation between the Amur leopards and the African leopards. *f3*- statistics were calculated using admixtools v7.0.2 (Patterson et al., 2012) with the haploid panel. We also used LocalNgsRelate (Severson et al., 2021) to infer identity-by-descent (IBD) sharing along the genome between the Zanzibar and other African leopards with default parameters.

### Admixture graph modelling

To model the ancestry of the Zanzibar leopard and other African leopards, we built admixture graphs using qpGraph (Patterson et al., 2012) with the haploid panel. First, we identified samples with the same geographical origin that can be grouped into one population with qpWave (Patterson et al., 2012). qpWave tests whether the samples in one population (termed as the left population) have ancestry of N independent migrations from another list of reference populations (termed the right population). A list of African leopard samples sharing the same geographical origin was put as the left population, and the rest of the African leopard samples and the Amur leopard were used as the right population. We removed a sample each time from the left population until the remaining samples were tested as deriving from a single migration from the right population, and hence grouped into one population (Table. S1). Then, we built a base graph using the Amur leopards as the outgroup, and four African leopard populations including Ghana, Zambia, Namibia, and East Tanzania (TanzaniaE). qpBrute (Leathlobhair et al., 2018; Liu et al., 2019) was used to perform a heuristic search in the graph space, and we identified the best-fitted graph with minimum admixture event and lowest z-score (Fig. S4). The Zanzibar leopard and one of the remaining African leopard populations were added separately to the based graph with qpBrute, in order to find the best-fitted admixture graph for each pair of populations.

### Heterozygosity, Runs of Homozygosity and Genetic Load estimation

We estimated heterozygosity for each sample using ANGSD. The folded site frequency spectrum (SFS) was first estimated using the domestic cat reference genome as the ancestral state. Then heterozygosity was calculated based on the SFS. We excluded transversion sites for the SFS estimation. Only samples that had sequence depth above 8× and that passed the quality control thresholds were included in this analysis. To infer runs of homozygosity within leopard genomes, we used Plink v1.9 with the following settings to adjust for the possible short fragment from ancient DNA, and allow more missing in each ROH region. The detailed parameters were *--homozyg-snp 30 --homozyg-kb 500 --homozyg-density 30 --homozyg-gap 1000 --homozyg-window-snp 30 --homozyg-window-het 1 --homozyg-window-missing 10 -- homozyg-window-threshold 0.05*.

Genetic load for leopards were estimated with two different approaches. Genetic load can be quantified by measuring the accumulation of deleterious mutations in the genome. We identified deleterious mutations with two different approaches. The first method considers deleterious mutations as homozygous derived mutations in leopards using conservation scores based on multi-species alignment. PhyloP and PhastCons scores based on 100 vertebrate species were used (Hubisz et al., 2011). The conservation scores were stored with reference to the hg38 human reference genome. We retained the loci with the top 5 percentage score, and mapped the coordinates in hg38 to felcat9 using *liftover*. The domestic cat reference genome was used as the ancestral state for the leopard genomes. Genetic load was calculated as the sum of the conservation score of homozygous derived loci in each individual. In order to reduce the effect of ancient DNA, we excluded transitions.

The other method considered deleterious mutations as mutations to have functional effects in coding regions. We annotated variable sites in leopards using SnpEff v5.1 (Cingolani et al., 2012). Genetic load was quantified as the total count of homozygous derived loss of function and nonsynonymous mutations/synonymous mutations in each individual.

### Gene Ontology for genes with homozygous derived mutations specific to the Zanzibar leopard

We generated the gene list containing homozygous derived mutations specific to the Zanzibar leopard sample. GO term and KEGG pathway enrichment analyses were performed with the KOBAS 3.0 web server (Bu et al., 2021). A threshold of P-value<0.05 after multiple testing corrections via FDR estimation was considered significant.

## Data and code availability

Whole-genome re-sequencing data are deposited at NCBI GenBank under the accession number of PRJNA892480. The detailed data processing pipeline for this project is available at https://github.com/xin-sun-popgen/zanzibar_leopard.

## Supplemental information titles and legends

**Figure S1. Maximum likelihood phylogeny of mitochondrial sequences.** Regions containing potential *numts* were excluded. Bootstrap support was labelled for major clades. The tiger mitochondrial sequence was used to root the phylogenetic tree.

**Figure S2. MDS analysis of all leopards included in the dataset.** Icon colours and shapes are labelled according to their geographical origin.

**Figure S3. Lack of IBD sharing between Zanzibar leopard and other African leopards.** As a comparison, IBD sharing between other African leopards was shown. Y axis refers to the IBD sharing status between the two individuals with 0 as no IBD, 1 as half IBD and 2 as full IBD sharing.

**Figure S4. Best-fitted admixture graphs modelling the ancestry of African leopards.** The Zanzibar leopard and one other African leopard were fitted into the starting graph consisting of major African leopard populations, including NW (northwest, Africa_Gha), Tanzania (TanzaniaE), Zambia, Namibia, using the Amur leopard (*P. p. orientalis*) as the outgroup.

**Table S1. Sequencing results of leopard specimens in this study**

**Table S2. Gene ontology and KEGG enrichment result of Zanzibar leopard unique homozygous SNPs**

**Table S3. Candidate gene list for body size in canid and coat colour pattern in domestic cats**

**Table S4. Body size and coat colour pattern related genes containing homozygous derived alleles in the Zanzibar leopards**

## References

Athreya, V., Odden, M., Linnell, J.D.C., Krishnaswamy, J., Ullas Karanth, K., 2016. A cat among the dogs: leopard Panthera pardus diet in a human-dominated landscape in western Maharashtra, India. Oryx 50, 156–162.

Bu, D., Luo, H., Huo, P., Wang, Z., Zhang, S., He, Z., Wu, Y., Zhao, L., Liu, J., Guo, J., Fang, S., Cao, W., Yi, L., Zhao, Y., Kong, L., 2021. KOBAS-i: intelligent prioritization and exploratory visualization of biological functions for gene enrichment analysis. Nucleic Acids Res. 49, W317–W325.

Butchart, S.H.M., Walpole, M., Collen, B., van Strien, A., Scharlemann, J.P.W., Almond, R.E.A., Baillie, J.E.M., Bomhard, B., Brown, C., Bruno, J., Carpenter, K.E., Carr, G.M., Chanson, J., Chenery, A.M., Csirke, J., Davidson, N.C., Dentener, F., Foster, M., Galli, A., Galloway, J.N., Genovesi, P., Gregory, R.D., Hockings, M., Kapos, V., Lamarque, J.-F., Leverington, F., Loh, J., McGeoch, M.A., McRae, L., Minasyan, A., Hernández Morcillo, M., Oldfield, T.E.E., Pauly, D., Quader, S., Revenga, C., Sauer, J.R., Skolnik, B., Spear, D., Stanwell-Smith, D., Stuart, S.N., Symes, A., Tierney, M., Tyrrell, T.D., Vié, J.-C., Watson, R., 2010. Global biodiversity: indicators of recent declines. Science 328, 1164–1168.

Chang, C.C., Chow, C.C., Tellier, L.C., Vattikuti, S., Purcell, S.M., Lee, J.J., 2015. Second-generation PLINK: rising to the challenge of larger and richer datasets. Gigascience 4, 7.

Chase, K., Carrier, D.R., Adler, F.R., Jarvik, T., Ostrander, E.A., Lorentzen, T.D., Lark, K.G., 2002. Genetic basis for systems of skeletal quantitative traits: principal component analysis of the canid skeleton. Proc. Natl. Acad. Sci. U. S. A. 99, 9930–9935.

Cingolani, P., Platts, A., Wang, L.L., Coon, M., Nguyen, T., Wang, L., Land, S.J., Lu, X., Ruden, D.M., 2012. A program for annotating and predicting the effects of single nucleotide polymorphisms, SnpEff: SNPs in the genome of Drosophila melanogaster strain w1118; iso-2; iso-3. Fly 6, 80–92.

Colossal^TM^/ A New Dawn of Genetics [WWW Document], n.d. . Colossal. URL https://colossal.com/

Dabney, J., Knapp, M., Glocke, I., Gansauge, M.-T., Weihmann, A., Nickel, B., Valdiosera, C., García, N., Pääbo, S., Arsuaga, J.-L., Meyer, M., 2013. Complete mitochondrial genome sequence of a Middle Pleistocene cave bear reconstructed from ultrashort DNA fragments. Proc. Natl. Acad. Sci. U. S. A. 110, 15758–15763.

Eizirik, E., David, V.A., Buckley-beason, V., Roelke, M.E., Scha, A.A., Hannah, S.S., Narfstro, K., Brien, S.J.O., Menotti-raymond, M., 2010. Defining and Mapping Mammalian Coat Pattern Genes: Multiple Genomic Regions Implicated in Domestic Cat Stripes and Spots. https://doi.org/10.1534/genetics.109.109629

Eizirik, E., Yuhki, N., Johnson, W.E., Menotti-Raymond, M., Hannah, S.S., O’Brien, S.J., 2003. Molecular genetics and evolution of melanism in the cat family. Curr. Biol. 13, 448–453.

Fernandez, F.A.S., Rheingantz, M.L., Genes, L., Kenup, C.F., Galliez, M., Cezimbra, T., Cid, B., Macedo, L., Araujo, B.B.A., Moraes, B.S., Monjeau, A., Pires, A.S., 2017. Rewilding the Atlantic Forest: Restoring the fauna and ecological interactions of a protected area. Perspectives in Ecology and Conservation 15, 308–314.

Figueiró, H.V., Li, G., Trindade, F.J., Assis, J., Pais, F., Fernandes, G., Santos, S.H.D., Hughes, G.M., Komissarov, A., Antunes, A., Trinca, C.S., Rodrigues, M.R., Linderoth, T., Bi, K., Silveira, L., Azevedo, F.C.C., Kantek, D., Ramalho, E., Brassaloti, R.A., Villela, P.M.S., Nunes, A.L.V., Teixeira, R.H.F., Morato, R.G., Loska, D., Saragüeta, P., Gabaldón, T., Teeling, E.C., O’Brien, S.J., Nielsen, R., Coutinho, L.L., Oliveira, G., Murphy, W.J., Eizirik, E., 2017. Genome-wide signatures of complex introgression and adaptive evolution in the big cats. Science Advances 3, e1700299.

Frankham, R., 1997. Do island populations have less genetic variation than mainland populations? Heredity 78 ( Pt 3), 311–327.

Gansauge, M.-T., Meyer, M., 2013. Single-stranded DNA library preparation for the sequencing of ancient or damaged DNA. Nat. Protoc. 8, 737–748.

Germano, J.M., Bishop, P.J., 2009. Suitability of amphibians and reptiles for translocation. Conserv. Biol. 23, 7–15.

Goldman, H.V., Walsh, M.T., 2002. Is The Zanzibar Leopard (*Panthera pardus adersi*) Extinct. J. East Afr. Nat. Hist. Soc. Natl. Mus. 91, 15–25.

Griffiths, C.J., Zuël, N., Jones, C.G., Ahamud, Z., Harris, S., 2013. Assessing the potential to restore historic grazing ecosystems with tortoise ecological replacements. Conserv. Biol. 27, 690–700.

Havmøller, R.W., Jacobsen, N.S., Havmøller, L.W., Rovero, F., Scharff, N., Bohmann, K., 2021. DNA metabarcoding reveals that African leopard diet varies between habitats. Afr. J. Ecol. 59, 37–50.

Hayward, J.J., Castelhano, M.G., Oliveira, K.C., Corey, E., Balkman, C., Baxter, T.L., Casal, M.L., Center, S.A., Fang, M., Garrison, S.J., Kalla, S.E., Korniliev, P., Kotlikoff, M.I., Moise, N.S., Shannon, L.M., Simpson, K.W., Sutter, N.B., Todhunter, R.J., Boyko, A.R., 2016. Complex disease and phenotype mapping in the domestic dog. Nat. Commun. 7, 10460.

Hoopes, B.C., Rimbault, M., Liebers, D., Ostrander, E.A., Sutter, N.B., 2012. The insulin-like growth factor 1 receptor (IGF1R) contributes to reduced size in dogs. Mamm. Genome 23, 780–790.

Hubisz, M.J., Pollard, K.S., Siepel, A., 2011. PHAST and RPHAST: phylogenetic analysis with space/time models. Brief. Bioinform. 12, 41–51.

Hunter, E.A., Gibbs, J.P., Cayot, L.J., Tapia, W., 2013. Equivalency of Galápagos giant tortoises used as ecological replacement species to restore ecosystem functions. Conserv. Biol. 27, 701–709.

Iacona, G., Maloney, R.F., Chadès, I., Bennett, J.R., Seddon, P.J., Possingham, H.P., 2017. Prioritizing revived species: what are the conservation management implications of de-extinction? Funct. Ecol. 31, 1041–1048.

IUCN/SSC, 2016. IUCN SSC Guiding principles on Creating Proxies of Extinct Species for Conservation Benefit. Version 1.0. Gland, Switzerland: IUCN Species Survival Commission.

Jones, P., Chase, K., Martin, A., Davern, P., Ostrander, E.A., Lark, K.G., 2008. Single-nucleotide-polymorphism-based association mapping of dog stereotypes. Genetics 179, 1033–1044.

Kaelin, C.B., McGowan, K.A., Barsh, G.S., 2021. Developmental genetics of color pattern establishment in cats. Nat. Commun. 12, 5127.

Kapp, J.D., Green, R.E., Shapiro, B., 2021. A Fast and Efficient Single-stranded Genomic Library Preparation Method Optimized for Ancient DNA. J. Hered. 112, 241–249.

Keedwell, R.J., Maloney, R.F., Murray, D.P., 2002. Predator control for protecting kaki (*Himantopus novaezelandiae*)—lessons from 20 years of management. Biol. Conserv. 105, 369–374.

Korneliussen, T.S., Albrechtsen, A., Nielsen, R., 2014. ANGSD: analysis of next generation sequencing data. BMC Bioinformatics 15, 356.

Larison, B., Kaelin, C.B., Harrigan, R., Henegar, C., Rubenstein, D.I., Kamath, P., Aschenborn, O., Smith, T.B., Barsh, G.S., 2021. Population structure, inbreeding and stripe pattern abnormalities in plains zebras. Mol. Ecol. 30, 379–390.

Leathlobhair, M.N., Perri, A.R., Irving-Pease, E.K., Witt, K.E., Linderholm, A., Haile, J., Lebrasseur, O., Ameen, C., Blick, J., Boyko, A.R., Brace, S., Cortes, Y.N., Crockford, S.J., Devault, A., Dimopoulos, E.A., Eldridge, M., Enk, J., Gopalakrishnan, S., Gori, K., Grimes, V., Guiry, E., Hansen, A.J., Hulme-Beaman, A., Johnson, J., Kitchen, A., Kasparov, A.K., Kwon, Y.-M., Nikolskiy, P.A., Lope, C.P., Manin, A., Martin, T., Meyer, M., Myers, K.N., Omura, M., Rouillard, J.-M., Pavlova, E.Y., Sciulli, P., Sinding, M.-H.S., Strakova, A., Ivanova, V.V., Widga, C., Willerslev, E., Pitulko, V.V., Barnes, I., Gilbert, M.T.P., Dobney, K.M., Malhi, R.S., Murchison, E.P., Larson, G., Frantz, L.A.F., 2018. The evolutionary history of dogs in the Americas. Science 361, 81– 85.

Li, H., Handsaker, B., Wysoker, A., Fennell, T., Ruan, J., Homer, N., Marth, G., Abecasis, G., Durbin, R., 2009. The sequence alignment/map format and SAMtools. Bioinformatics 25, 2078–2079.

Lin, J., Duchêne, D., Carøe, C., Smith, O., Ciucani, M.M., Niemann, J., Richmond, D., Greenwood, A.D., MacPhee, R., Zhang, G., Gopalakrishnan, S., Gilbert, M.T.P., 2022. Probing the genomic limits of de-extinction in the Christmas Island rat. Curr. Biol. 0. https://doi.org/10.1016/j.cub.2022.02.027

Liu, L., Bosse, M., Megens, H.-J., Frantz, L.A.F., Lee, Y.-L., Irving-Pease, E.K., Narayan, G., Groenen, M.A.M., Madsen, O., 2019. Genomic analysis on pygmy hog reveals extensive interbreeding during wild boar expansion. Nat. Commun. 10, 1992.

Lorimer, J., Sandom, C., Jepson, P., Doughty, C., Barua, M., Kirby, K.J., 2015. Rewilding: Science, Practice, and Politics. Annu. Rev. Environ. Resour. 40, 39–62.

Mak, S.S.T., Gopalakrishnan, S., Carøe, C., Geng, C., Liu, S., Sinding, M.-H.S., Kuderna, L.F.K., Zhang, W., Fu, S., Vieira, F.G., Germonpré, M., Bocherens, H., Fedorov, S., Petersen, B., Sicheritz-Pontén, T., Marques-Bonet, T., Zhang, G., Jiang, H., Gilbert, M.T.P., 2017. Comparative performance of the BGISEQ-500 vs Illumina HiSeq2500 sequencing platforms for palaeogenomic sequencing. Gigascience 6, 1–13.

Meisner, J., Albrechtsen, A., 2018. Inferring Population Structure and Admixture Proportions in Low-Depth NGS Data. Genetics 210, 719–731.

Meyer, M., Kircher, M., 2010. Illumina Sequencing Library Preparation for Highly Multiplexed Target Capture and Sequencing. Cold Spring Harb. Protoc. 2010, db.prot5448.

Novak, B.J., 2018. De-Extinction. Genes 9. https://doi.org/10.3390/genes9110548

Paijmans, J.L.A., Barlow, A., Becker, M.S., Cahill, J.A., Fickel, J., Förster, D.W.G., Gries, K., Hartmann, S., Havmøller, R.W., Henneberger, K., Kern, C., Kitchener, A.C., Lorenzen, E.D., Mayer, F., OBrien, S.J., von Seth, J., Sinding, M.-H.S., Spong, G., Uphyrkina, O., Wachter, B., Westbury, M.V., Dalén, L., Bhak, J., Manica, A., Hofreiter, M., 2021. African and Asian leopards are highly differentiated at the genomic level. Curr. Biol. 31, 1872–1882.e5.

Pakenham, R.H.W., 1984. The mammals of Zanzibar and Pemba islands. Pakenham.

Parker, H.G., VonHoldt, B.M., Quignon, P., Margulies, E.H., Shao, S., Mosher, D.S., Spady, T.C., Elkahloun, A., Cargill, M., Jones, P.G., Maslen, C.L., Acland, G.M., Sutter, N.B., Kuroki, K., Bustamante, C.D., Wayne, R.K., Ostrander, E.A., 2009. An expressed fgf4 retrogene is associated with breed-defining chondrodysplasia in domestic dogs. Science 325, 995–998.

Park, S.D.E., Magee, D.A., McGettigan, P.A., Teasdale, M.D., Edwards, C.J., Lohan, A.J., Murphy, A., Braud, M., Donoghue, M.T., Liu, Y., Chamberlain, A.T., Rue-Albrecht, K., Schroeder, S., Spillane, C., Tai, S., Bradley, D.G., Sonstegard, T.S., Loftus, B.J., Machugh, D.E., 2015. Genome sequencing of the extinct Eurasian wild aurochs, Bos primigenius, illuminates the phylogeography and evolution of cattle. Genome Biol. 1–15.

Patterson, N., Moorjani, P., Luo, Y., Mallick, S., Rohland, N., Zhan, Y., Genschoreck, T., Webster, T., Reich, D., 2012. Ancient admixture in human history. Genetics 192, 1065– 1093.

Pečnerová, P., Garcia-Erill, G., Liu, X., Nursyifa, C., Waples, R.K., Santander, C.G., Quinn, L., Frandsen, P., Meisner, J., Stæger, F.F., Rasmussen, M.S., Brüniche-Olsen, A., Hviid Friis Jørgensen, C., da Fonseca, R.R., Siegismund, H.R., Albrechtsen, A., Heller, R., Moltke, I., Hanghøj, K., 2021. High genetic diversity and low differentiation reflect the ecological versatility of the African leopard. Curr. Biol. 31, 1862–1871.e5.

Pereira, H.M., Leadley, P.W., Proença, V., Alkemade, R., Scharlemann, J.P.W., Fernandez-Manjarrés, J.F., Araújo, M.B., Balvanera, P., Biggs, R., Cheung, W.W.L., Chini, L., Cooper, H.D., Gilman, E.L., Guénette, S., Hurtt, G.C., Huntington, H.P., Mace, G.M., Oberdorff, T., Revenga, C., Rodrigues, P., Scholes, R.J., Sumaila, U.R., Walpole, M., 2010. Scenarios for global biodiversity in the 21st century. Science 330, 1496–1501.

Perino, A., Pereira, H.M., Navarro, L.M., Fernández, N., Bullock, J.M., Ceaușu, S., Cortés-Avizanda, A., van Klink, R., Kuemmerle, T., Lomba, A., Pe’er, G., Plieninger, T., Rey Benayas, J.M., Sandom, C.J., Svenning, J.-C., Wheeler, H.C., 2019. Rewilding complex ecosystems. Science 364. https://doi.org/10.1126/science.aav5570

Peterschmitt, M., Grain, F., Arnaud, B., Deléage, G., Lambert, V., 2009. Mutation in the melanocortin 1 receptor is associated with amber colour in the Norwegian Forest Cat. Anim. Genet. 40, 547–552.

Plassais, J., Kim, J., Davis, B.W., Karyadi, D.M., Hogan, A.N., Harris, A.C., Decker, B., Parker, H.G., Ostrander, E.A., 2019. Whole genome sequencing of canids reveals genomic regions under selection and variants influencing morphology. Nat. Commun. 10, 1489.

Plassais, J., vonHoldt, B.M., Parker, H.G., Carmagnini, A., Dubos, N., Papa, I., Bevant, K., Derrien, T., Hennelly, L.M., Whitaker, D.T., Harris, A.C., Hogan, A.N., Huson, H.J., Zaibert, V.F., Linderholm, A., Haile, J., Fest, T., Habib, B., Sacks, B.N., Benecke, N., Outram, A.K., Sablin, M.V., Germonpré, M., Larson, G., Frantz, L., Ostrander, E.A., 2022. Natural and human-driven selection of a single non-coding body size variant in ancient and modern canids. Curr. Biol. 32, 889–897.e9.

Prendergast, M.E., Rouby, H., Punnwong, P., Marchant, R., Crowther, A., Kourampas, N., Shipton, C., Walsh, M., Lambeck, K., Boivin, N.L., 2016. Continental Island Formation and the Archaeology of Defaunation on Zanzibar, Eastern Africa. PLoS One 11, e0149565.

Ramos-Madrigal, J., Sinding, M.-H.S., Carøe, C., Mak, S.S.T., Niemann, J., Samaniego Castruita, J.A., Fedorov, S., Kandyba, A., Germonpré, M., Bocherens, H., Feuerborn, T.R., Pitulko, V.V., Pavlova, E.Y., Nikolskiy, P.A., Kasparov, A.K., Ivanova, V.V., Larson, G., Frantz, L.A.F., Willerslev, E., Meldgaard, M., Petersen, B., Sicheritz-Ponten, T., Bachmann, L., Wiig, Ø., Hansen, A.J., Gilbert, M.T.P., Gopalakrishnan, S., 2021. Genomes of Pleistocene Siberian Wolves Uncover Multiple Extinct Wolf Lineages. Curr. Biol. 31, 198–206.e8.

Richmond, D.J., Sinding, M.-H.S., Gilbert, M.T.P., 2016. The potential and pitfalls of de-extinction. Zool. Scr. 45, 22–36.

Rimbault, M., Beale, H.C., Schoenebeck, J.J., Hoopes, B.C., Allen, J.J., Kilroy-Glynn, P., Wayne, R.K., Sutter, N.B., Ostrander, E.A., 2013. Derived variants at six genes explain nearly half of size reduction in dog breeds. Genome Res. 23, 1985–1995.

Ripple, W.J., Beschta, R.L., 2012. Trophic cascades in Yellowstone: The first 15 years after wolf reintroduction. Biol. Conserv. 145, 205–213.

Schubert, M., Ermini, L., Der Sarkissian, C., Jónsson, H., Ginolhac, A., Schaefer, R., Martin, M.D., Fernández, R., Kircher, M., McCue, M., Willerslev, E., Orlando, L., 2014. Characterization of ancient and modern genomes by SNP detection and phylogenomic and metagenomic analysis using PALEOMIX. Nat. Protoc. 9, 1056–1082.

Seddon, P.J., Griffiths, C.J., Soorae, P.S., Armstrong, D.P., 2014a. Reversing defaunation: restoring species in a changing world. Science 345, 406–412.

Seddon, P.J., Moehrenschlager, A., Ewen, J., 2014b. Reintroducing resurrected species: selecting DeExtinction candidates. Trends Ecol. Evol. 29, 140–147.

Severson, A.L., Korneliussen, T.S., Moltke, I., 2021. LocalNgsRelate: a software tool for inferring IBD sharing along the genome between pairs of individuals from low-depth NGS data. Bioinformatics. https://doi.org/10.1093/bioinformatics/btab732

Shapiro, B., 2017. Pathways to de-extinction: how close can we get to resurrection of an extinct species? Funct. Ecol. 31, 996–1002.

Shapiro, B., 2015. How to Clone a Mammoth. Princeton University Press, Princeton, NJ. https://doi.org/10.1515/9781400865482

Sinding, M.-H.S., Gilbert, M.T.P., 2016. The Draft Genome of Extinct European Aurochs and its Implications for De-Extinction. Open Quaternary 2, e5753.

Skotte, L., Korneliussen, T.S., Albrechtsen, A., 2013. Estimating individual admixture proportions from next generation sequencing data. Genetics 195, 693–702.

Stokstad, E., 2015. Bringing back the aurochs. Science 350, 1144–1147.

Sutter, N.B., Bustamante, C.D., Chase, K., Gray, M.M., Zhao, K., Zhu, L., Padhukasahasram, B., Karlins, E., Davis, S., Jones, P.G., Quignon, P., Johnson, G.S., Parker, H.G., Fretwell, N., Mosher, D.S., Lawler, D.F., Satyaraj, E., Nordborg, M., Lark, K.G., Wayne, R.K., Ostrander, E.A., 2007. A single IGF1 allele is a major determinant of small size in dogs. Science 316, 112–115.

Svenning, J.-C., Pedersen, P.B.M., Donlan, C.J., Ejrnæs, R., Faurby, S., Galetti, M., Hansen, D.M., Sandel, B., Sandom, C.J., Terborgh, J.W., Vera, F.W.M., 2016. Science for a wilder Anthropocene: Synthesis and future directions for trophic rewilding research. Proc. Natl. Acad. Sci. U. S. A. 113, 898–906.

The Quagga Project [WWW Document], n.d. . Quaggaproject. URL https://www.quaggaproject.org/

The Tauros programme [WWW Document], n.d. . TaurOs Project. URL http://www.taurosproject.com/

Uphyrkina, O., Miquelle, D., Quigley, H., Driscoll, C., O’Brien, S.J., 2002. Conservation genetics of the Far Eastern leopard (Panthera pardus orientalis). J. Hered. 93, 303–311.

Vaysse, A., Ratnakumar, A., Derrien, T., Axelsson, E., Rosengren Pielberg, G., Sigurdsson, S., Fall, T., Seppälä, E.H., Hansen, M.S.T., Lawley, C.T., Karlsson, E.K., LUPA Consortium, Bannasch, D., Vilà, C., Lohi, H., Galibert, F., Fredholm, M., Häggström, J., Hedhammar, A., André, C., Lindblad-Toh, K., Hitte, C., Webster, M.T., 2011. Identification of genomic regions associated with phenotypic variation between dog breeds using selection mapping. PLoS Genet. 7, e1002316.

Walsh, M.T., Goldman, H.V., 2008. Updating the inventory of Zanzibar leopard specimens. Cat News 49, 4–6.

Walsh, M.T., Goldman, H.V., 2007. Killing the king: the demonization and extermination of the Zanzibar leopard, in: Dounias, E., Motte-Florac, M.E., Dunham, M. (Eds.), Animal Symbolism: Animals, Keystone of the Relationship between Man and Nature? IRD, pp. 1133–1182.

