## Supplemental tables and figures for "A genomic exploration of the possible de-extirpation of the Zanzibar leopard"

### Supplementary Materials

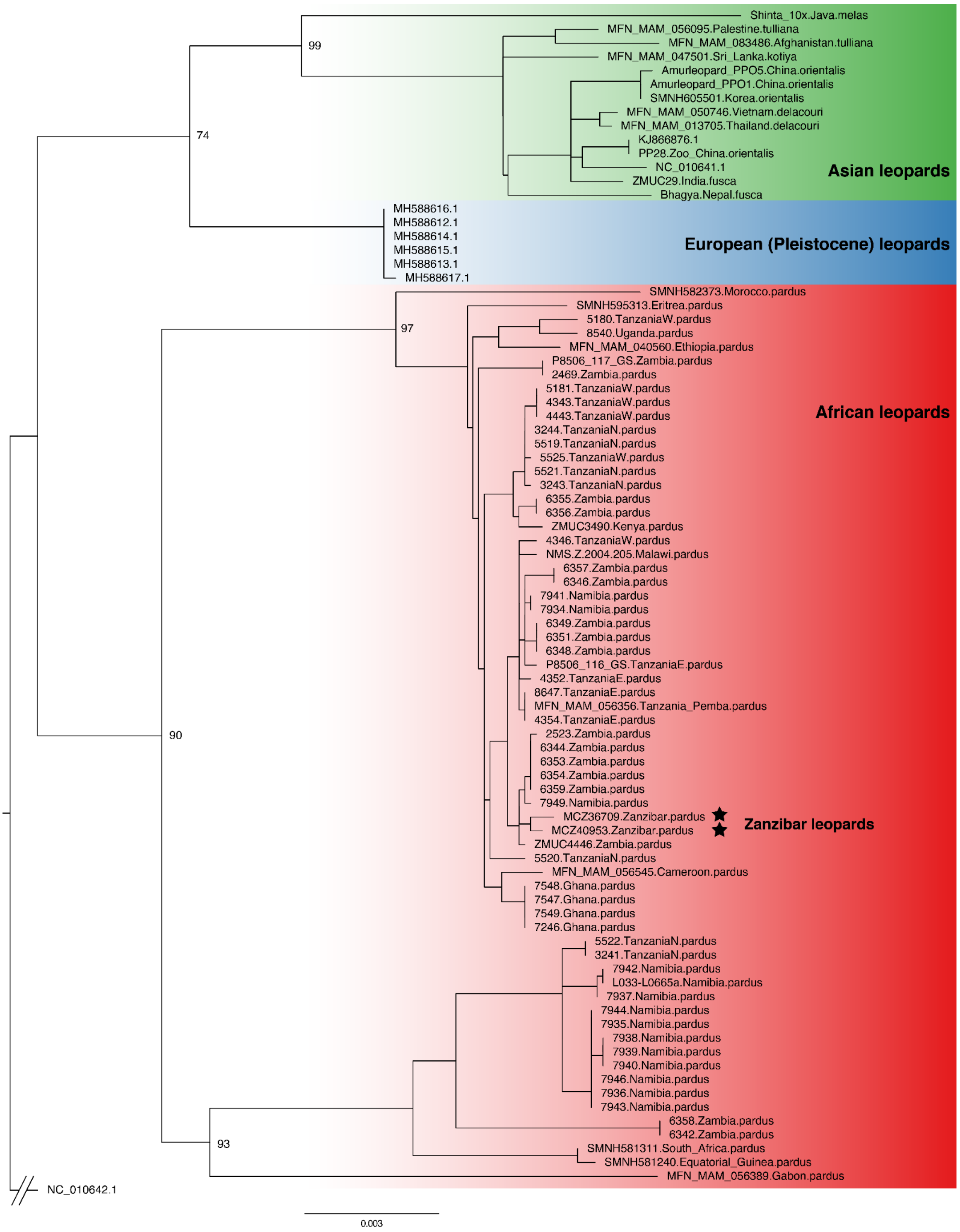


**Figure S1. Maximum likelihood phylogeny of mitochondrial sequences.** Regions containing potential *numts* were excluded. Bootstrap support was labelled for major clades. The tiger mitochondrial sequence was used to root the phylogenetic tree.

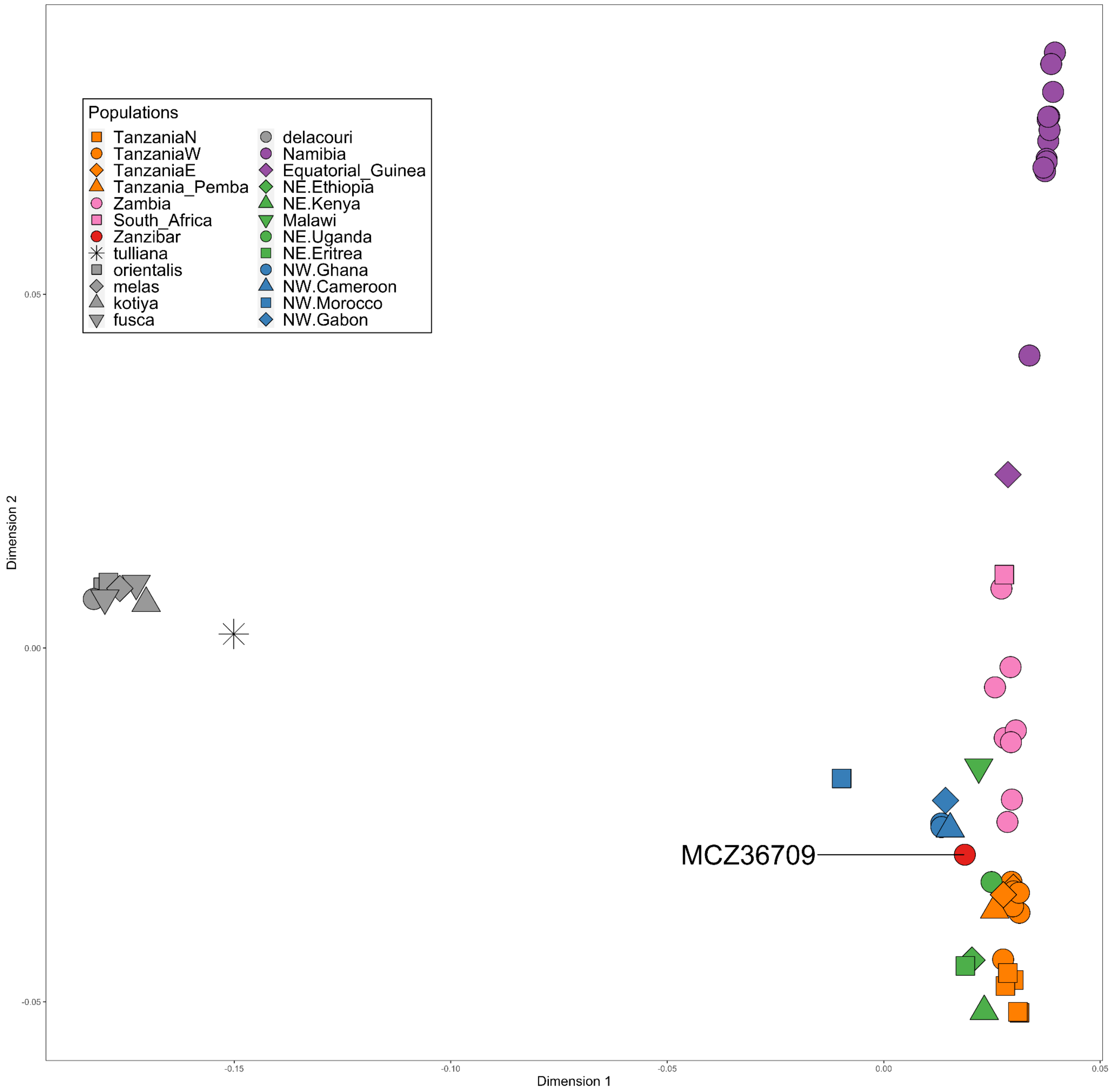


**Figure S2. MDS analysis of all leopards included in the dataset.** Icon colours and shapes are labelled according to their geographical origin.

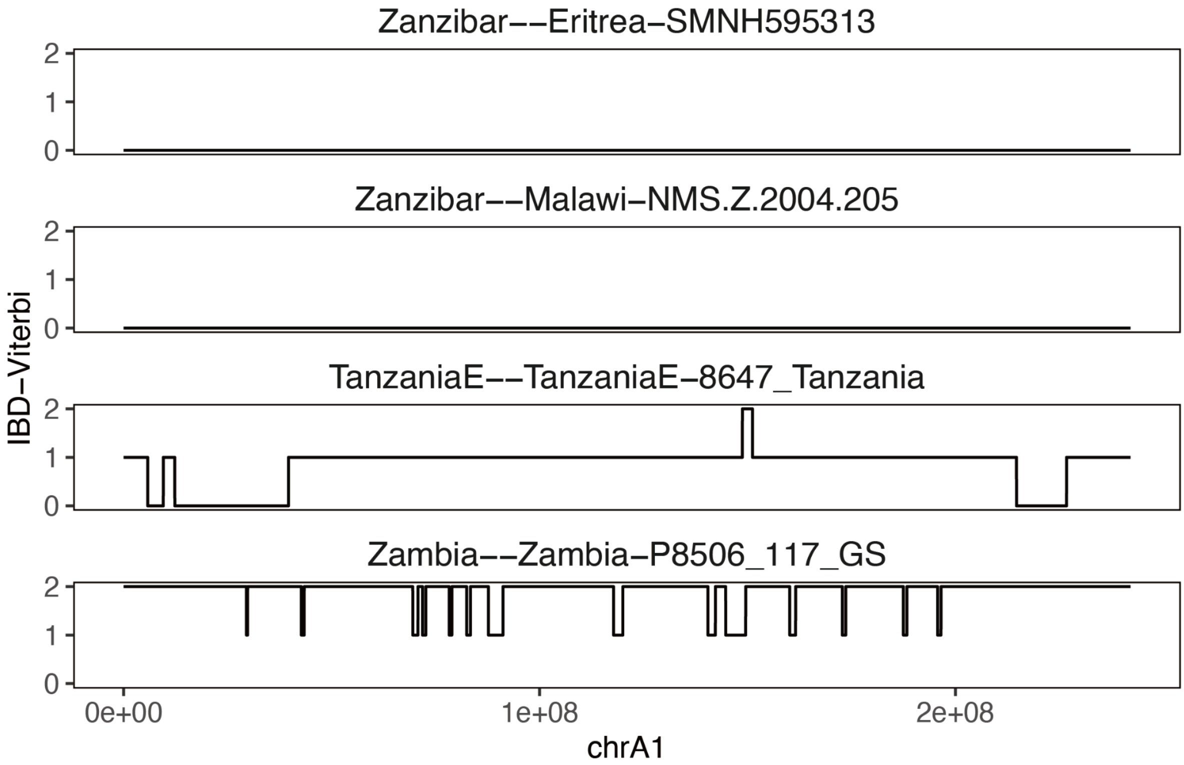


**Figure S3. Lack of IBD sharing between Zanzibar leopard and other African leopards.** As a comparison, IBD sharing between other African leopards was shown. Y axis refers to the IBD sharing status between the two individuals with 0 as no IBD, 1 as half IBD and 2 as full IBD sharing.

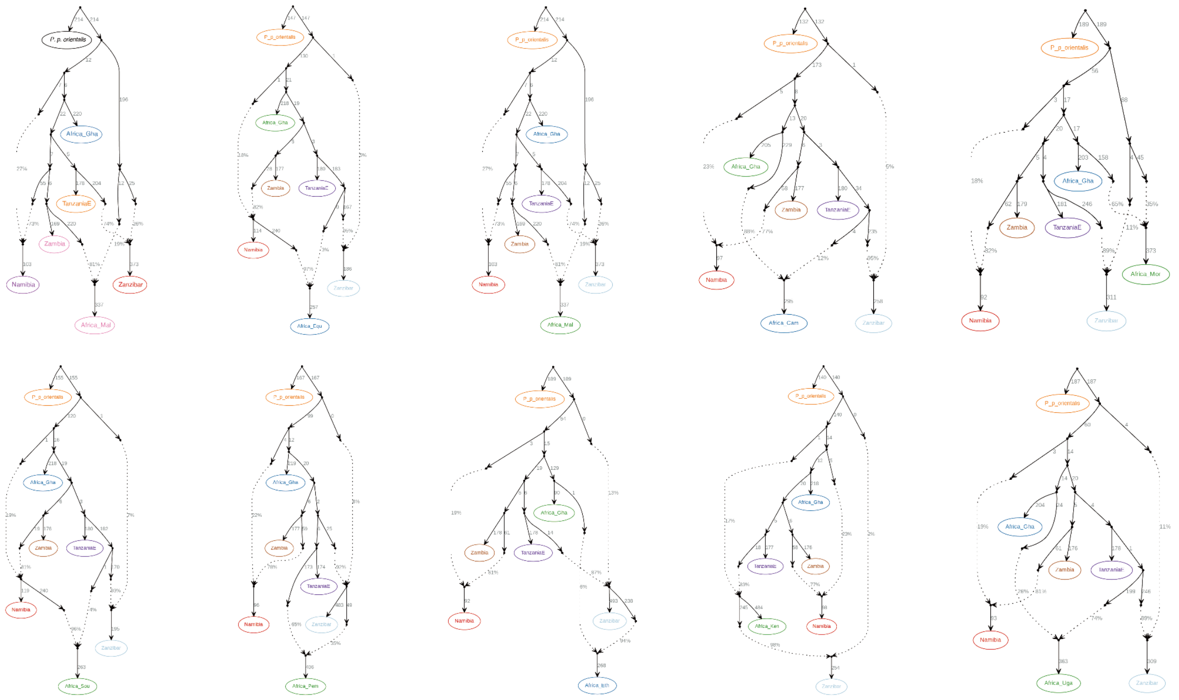

**Figure S4. Best-fitted admixture graphs modelling the ancestry of African leopards.** The Zanzibar leopard and one other African leopard were fitted into the starting graph consisting of major African leopard populations, including NW (northwest, Africa_Gha), Tanzania (TanzaniaE), Zambia, Namibia, using the Amur leopard (*P. p. orientalis*) as the outgroup.

### Table S1. Sequencing results of leopard specimens in this study

| Sample ID | Common name | Scientific name | Population assigned | Relatedness filter | Error rate filter | Sequencing depth (X) | Sequencing depth after filtering (X) | Sex | Pseudohaploid panel | Genotype calling (samtools) | PCA/ MDS | Tree mix | Admixture Graph group | qpWave group | PSMC | Sample/Date source |
|---|---|---|---|---|---|---|---|---|---|---|---|---|---|---|---|---|
| MCZ36709 | Zanzibar leopard | *P. p. pardus* | Zanzibar | No | No | 38.4 | 30.4 | M | Yes | Yes | Yes | Yes | Zanzibar | Zanzibar | Yes | MCZ voucher numbers 36709, Harvard Museum of Comparative Zoology |
| MCZ40953 | Zanzibar leopard | *P. p. pardus* | Zanzibar | | | 0.40 | exclude | | | | | | Exclude | Exclude | | MCZ voucher numbers 40953, Harvard Museum of Comparative Zoology |
| Z1209 | Zanzibar leopard | *P. p. pardus* | Zanzibar | | | 0.00 | exclude | | | | | | Exclude | Exclude | | the Zanzibar Museum (ID: Z 1209) |
| MFN_MAM_083486 | Persian leopard | *P. p. tulliana* | tulliana | No | Yes, transversion only | 11.7 | 7.3 | F | No | Yes | Yes | No | Exclude | Exclude | Yes | Paijmans *et al.* 2021 |
| MFN_MAM_056095 | Persian leopard | *P. p. tulliana* | tulliana | No | No | 5.7 | 3.9 | F | Yes | | Yes | Yes | Exclude | Exclude | | Paijmans *et al.* 2021 |
| L033-L0665a | African leopard | *P. p. pardus* | Namibia | No | No | 32.5 | 26.6 | M | Yes | Yes | Yes | Yes | Namibia | Namibia | Yes | Paijmans *et al.* 2021 |
| 7942_Namibia | African leopard | *P. p. pardus* | Namibia | No | No | 18.4 | 15.5 | M | Yes | Yes | Yes | Yes | Namibia | Namibia | | Patrícia *et al.* 2021 |
| ZMUC4446 | African leopard | *P. p. pardus* | Zambia | No | No | 17.9 | 14.7 | F | Yes | Yes | Yes | Yes | Zambia | Exclude | Yes | Paijmans *et al.* 2021 |
| 3241_Tanzania | African leopard | *P. p. pardus* | TanzaniaN | No | No | 17.8 | 14.4 | F | Yes | Yes | Yes | Yes | Tanzania NW | Exclude | Yes | Patrícia *et al.* 2021 |
| 4343_Tanzania | African leopard | *P. p. pardus* | TanzaniaW | No | No | 17.7 | 14.8 | M | Yes | Yes | Yes | Yes | Tanzania NW | Tanzania W | Yes | Patrícia *et al.* 2021 |
| 7547_Ghana | African leopard | *P. p. pardus* | NW.Ghana | No | No | 16.3 | 13.5 | F | Yes | Yes | Yes | Yes | Africa_Gha | Africa_Gha | Yes | Patrícia *et al.* 2021 |
| MFN_MAM_040560 | African leopard | *P. p. pardus* | NE.Ethiopia | No | No | 16.3 | 13.0 | M | Yes | Yes | Yes | Yes | Africa_NE | Africa_Eth | Yes | Paijmans *et al.* 2021 |

| | | | | | | | | | | | | | | | | |
|---|---|---|---|---|---|---|---|---|---|---|---|---|---|---|---|---|
| 4354_Tanzania | African leopard | *P. p. pardus* | TanzaniaE | No | No | 15.1 | 12.7 | M | Yes | Yes | Yes | Yes | TanzaniaE | TanzaniaE | Yes | Patrícia *et al.* 2021 |
| SMNH581311 | African leopard | *P. p. pardus* | South_Africa | No | No | 14.8 | 10.5 | F | Yes | Yes | Yes | Yes | Africa_S | Africa_Sou | Yes | Paijmans *et al.* 2021 |
| MFN_MAM_056356 | African leopard | *P. p. pardus* | Tanzania_Pemba | No | No | 14.4 | 11.5 | M | Yes | Yes | Yes | Yes | Africa_NE | Africa_Pem | | Paijmans *et al.* 2021 |
| ZMUC3490 | African leopard | *P. p. pardus* | NE.Kenya | No | No | 12.4 | 9.4 | F | Yes | Yes | Yes | Yes | Africa_NE | Africa_Ken | | Paijmans *et al.* 2021 |
| P8506_117_GS | African leopard | *P. p. pardus* | Zambia | No | No | 11.3 | 9.6 | M | Yes | Yes | Yes | Yes | Zambia | Exclude | | Paijmans *et al.* 2021 |
| MFN_MAM_056545 | African leopard | *P. p. pardus* | NW.Cameroon | No | No | 11.3 | 5.1 | M | Yes | Yes | Yes | Yes | Africa_Cam | Africa_Cam | | Paijmans *et al.* 2021 |
| P8506_116_GS | African leopard | *P. p. pardus* | TanzaniaE | No | No | 10.7 | 9.1 | F | Yes | Yes | Yes | Yes | TanzaniaE | Exclude | | Paijmans *et al.* 2021 |
| NMS.Z.2004.205 | African leopard | *P. p. pardus* | Malawi | No | No | 9.1 | 6.5 | F | Yes | Yes | Yes | Yes | Africa_NE | Africa_Mal | | Paijmans *et al.* 2021 |
| SMNH582373 | African leopard | *P. p. pardus* | NW.Morocco | No | No | 8.2 | 5.8 | F | Yes | Yes | Yes | Yes | Africa_Mor | Africa_Mor | | Paijmans *et al.* 2021 |
| SMNH581240 | African leopard | *P. p. pardus* | Equatorial_Guinea | No | No | 7.9 | 4.9 | M | Yes | | Yes | Yes | Africa_S | Africa_Equ | | Paijmans *et al.* 2021 |
| 7944_Namibia | African leopard | *P. p. pardus* | Namibia | No | No | 5.4 | 4.7 | M | Yes | | Yes | Yes | Namibia | Exclude | | Patrícia *et al.* 2021 |
| 6351_Zambia | African leopard | *P. p. pardus* | Zambia | No | No | 5.2 | 4.4 | F | Yes | | Yes | Yes | Zambia | Zambia | | Patrícia *et al.* 2021 |
| 4352_Tanzania | African leopard | *P. p. pardus* | TanzaniaE | No | No | 5.2 | 4.4 | M | Yes | | Yes | Yes | TanzaniaE | TanzaniaE | | Patrícia *et al.* 2021 |
| 7940_Namibia | African leopard | *P. p. pardus* | Namibia | No | No | 5.2 | 4.4 | F | Yes | | Yes | Yes | Namibia | Exclude | | Patrícia *et al.* 2021 |
| 5525_Tanzania | African leopard | *P. p. pardus* | TanzaniaW | No | No | 5.2 | 4.4 | M | Yes | | Yes | Yes | TanzaniaNW | Exclude | | Patrícia *et al.* 2021 |
| 6346_Zambia | African leopard | *P. p. pardus* | Zambia | No | No | 5.2 | 4.4 | F | Yes | | Yes | Yes | Zambia | Exclude | | Patrícia *et al.* 2021 |
| 7935_Namibia | African leopard | *P. p. pardus* | Namibia | No | No | 5.2 | 4.4 | M | Yes | | Yes | Yes | Namibia | Namibia | | Patrícia *et al.* 2021 |
| 6353_Zambia | African leopard | *P. p. pardus* | Zambia | No | No | 5.1 | 4.3 | F | Yes | | Yes | Yes | Zambia | Zambia | | Patrícia *et al.* 2021 |
| 7943_Namibia | African leopard | *P. p. pardus* | Namibia | Yes | No | 5.1 | 4.3 | M | No | | No | No | Exclude | Exclude | | Patrícia *et al.* 2021 |
| MFN_MAM_056389 | African leopard | *P. p. pardus* | NW.Gabon | No | No | 5.0 | 2.4 | M | Yes | | Yes | Yes | Exclude | Exclude | | Paijmans *et al.* 2021 |
| 7946_Namibia | African leopard | *P. p. pardus* | Namibia | No | No | 5.0 | 4.2 | M | Yes | | Yes | Yes | Namibia | Exclude | | Patrícia *et al.* 2021 |
| 7939_Namibia | African leopard | *P. p. pardus* | Namibia | No | No | 5.0 | 4.2 | M | Yes | | Yes | Yes | Namibia | Exclude | | Patrícia *et al.* 2021 |
| 6355_Zambia | African leopard | *P. p. pardus* | Zambia | No | No | 4.9 | 4.2 | M | Yes | | Yes | Yes | Zambia | Exclude | | Patrícia *et al.* 2021 |

| | | | | | | | | | | | | | | |
|---|---|---|---|---|---|---|---|---|---|---|---|---|---|---|
| 6344_Zambia | African leopard | *P. p. pardus* | Zambia | Yes | No | 4.9 | 4.1 | F | No | No | No | Exclude | Exclude | Patrícia *et al.* 2021 |
| 6357_Zambia | African leopard | *P. p. pardus* | Zambia | Yes | No | 4.8 | 4.1 | F | No | No | No | Exclude | Exclude | Patrícia *et al.* 2021 |
| 7937_Namibia | African leopard | *P. p. pardus* | Namibia | No | No | 4.8 | 4.1 | M | Yes | Yes | Yes | Namibia | Exclude | Patrícia *et al.* 2021 |
| 6349_Zambia | African leopard | *P. p. pardus* | Zambia | Yes | No | 4.8 | 4.0 | F | No | No | No | Exclude | Exclude | Patrícia *et al.* 2021 |
| 7949_Namibia | African leopard | *P. p. pardus* | Namibia | No | No | 4.7 | 4.0 | M | Yes | Yes | Yes | Namibia | Exclude | Patrícia *et al.* 2021 |
| 8540_Uganda | African leopard | *P. p. pardus* | NE.Uganda | No | No | 4.7 | 3.8 | M | Yes | Yes | Yes | Africa_Uga | Africa_Uga | Patrícia *et al.* 2021 |
| 7936_Namibia | African leopard | *P. p. pardus* | Namibia | No | No | 4.7 | 4.0 | F | Yes | Yes | Yes | Namibia | Exclude | Patrícia *et al.* 2021 |
| 7941_Namibia | African leopard | *P. p. pardus* | Namibia | No | No | 4.6 | 4.0 | M | Yes | Yes | Yes | Namibia | Exclude | Patrícia *et al.* 2021 |
| 7246_Ghana | African leopard | *P. p. pardus* | NW.Ghana | No | No | 4.6 | 3.8 | F | Yes | Yes | Yes | Africa_Gha | Africa_Gha | Patrícia *et al.* 2021 |
| 3244_Tanzania | African leopard | *P. p. pardus* | TanzaniaN | No | No | 4.6 | 3.8 | M | Yes | Yes | Yes | Tanzania NW | Tanzania N | Patrícia *et al.* 2021 |
| 6354_Zambia | African leopard | *P. p. pardus* | Zambia | Yes | No | 4.6 | 3.9 | F | No | No | No | Exclude | Exclude | Patrícia *et al.* 2021 |
| 7548_Ghana | African leopard | *P. p. pardus* | NW.Ghana | Yes | No | 4.6 | 3.8 | M | No | No | No | Exclude | Exclude | Patrícia *et al.* 2021 |
| 6342_Zambia | African leopard | *P. p. pardus* | Zambia | No | No | 4.6 | 3.7 | F | Yes | Yes | Yes | Zambia | Exclude | Patrícia *et al.* 2021 |
| 7466_Ghana | African leopard | *P. p. pardus* | NW.Ghana | No | Yes | 4.4 | 3.7 | F | No | No | No | Exclude | Exclude | Patrícia *et al.* 2021 |
| 3243_Tanzania | African leopard | *P. p. pardus* | TanzaniaN | Yes | No | 4.4 | 3.6 | F | No | No | No | Exclude | Exclude | Patrícia *et al.* 2021 |
| 2469_Zambia | African leopard | *P. p. pardus* | Zambia | Yes | No | 4.4 | 3.7 | M | No | No | No | Exclude | Exclude | Patrícia *et al.* 2021 |
| 6359_Zambia | African leopard | *P. p. pardus* | Zambia | Yes | No | 4.3 | 3.7 | F | No | No | No | Exclude | Exclude | Patrícia *et al.* 2021 |
| 7934_Namibia | African leopard | *P. p. pardus* | Namibia | No | No | 4.3 | 3.6 | F | Yes | Yes | Yes | Namibia | Exclude | Patrícia *et al.* 2021 |
| 5522_Tanzania | African leopard | *P. p. pardus* | TanzaniaN | No | No | 4.2 | 3.4 | M | Yes | Yes | Yes | Tanzania NW | Exclude | Patrícia *et al.* 2021 |
| 4443_Tanzania | African leopard | *P. p. pardus* | TanzaniaW | No | No | 4.2 | 3.5 | M | Yes | Yes | Yes | Tanzania NW | Exclude | Patrícia *et al.* 2021 |
| 5520_Tanzania | African leopard | *P. p. pardus* | TanzaniaN | No | No | 4.1 | 3.4 | M | Yes | Yes | Yes | Tanzania NW | Exclude | Patrícia *et al.* 2021 |
| 7938_Namibia | African leopard | *P. p. pardus* | Namibia | No | No | 4.1 | 3.5 | M | Yes | Yes | Yes | Namibia | Exclude | Patrícia *et al.* 2021 |
| 8647_Tanzania | African leopard | *P. p. pardus* | TanzaniaE | Yes | No | 4.1 | 3.2 | M | No | No | No | Exclude | Exclude | Patrícia *et al.* 2021 |

| | | | | | | | | | | | | | | | | |
|---|---|---|---|---|---|---|---|---|---|---|---|---|---|---|---|---|
| 4346_Tanzania | African leopard | *P. p. pardus* | TanzaniaW | No | No | 4.1 | 3.4 | M | Yes | | Yes | Yes | Tanzania NW | Exclude | | Patrícia *et al.* 2021 |
| 5181_Tanzania | African leopard | *P. p. pardus* | TanzaniaW | No | No | 3.9 | 3.2 | M | Yes | | Yes | Yes | Tanzania NW | Tanzania W | | Patrícia *et al.* 2021 |
| 6356_Zambia | African leopard | *P. p. pardus* | Zambia | Yes | No | 3.9 | 3.2 | M | No | | No | No | Exclude | Exclude | | Patrícia *et al.* 2021 |
| 5519_Tanzania | African leopard | *P. p. pardus* | TanzaniaN | No | No | 3.9 | 3.1 | M | Yes | | Yes | Yes | Tanzania NW | Tanzania N | | Patrícia *et al.* 2021 |
| 7549_Ghana | African leopard | *P. p. pardus* | NW.Ghana | Yes | No | 3.8 | 3.0 | F | No | | No | No | Exclude | Exclude | | Patrícia *et al.* 2021 |
| 5521_Tanzania | African leopard | *P. p. pardus* | TanzaniaN | Yes | No | 3.7 | 3.1 | M | No | | No | No | Exclude | Exclude | | Patrícia *et al.* 2021 |
| 7465_Ghana | African leopard | *P. p. pardus* | NW.Ghana | Yes | Yes | 3.7 | 3.1 | F | No | | No | No | Exclude | Exclude | | Patrícia *et al.* 2021 |
| 5180_Tanzania | African leopard | *P. p. pardus* | TanzaniaW | No | No | 3.7 | 2.9 | M | Yes | | Yes | Yes | Tanzania NW | Exclude | | Patrícia *et al.* 2021 |
| 6348_Zambia | African leopard | *P. p. pardus* | Zambia | Yes | No | 3.6 | 2.9 | F | No | | No | No | Exclude | Exclude | | Patrícia *et al.* 2021 |
| 6358_Zambia | African leopard | *P. p. pardus* | Zambia | Yes | No | 3.5 | 2.9 | F | No | | No | No | Exclude | Exclude | | Patrícia *et al.* 2021 |
| SMNH595313 | African leopard | *P. p. pardus* | NE.Eritrea | No | No | 3.5 | 2.6 | F | Yes | | Yes | Yes | Exclude | Exclude | | Paijmans *et al.* 2021 |
| 6350_Zambia | African leopard | *P. p. pardus* | Zambia | No | Yes | 3.4 | 0.6 | F | No | | No | No | Exclude | Exclude | | Patrícia *et al.* 2021 |
| 2523_Zambia | African leopard | *P. p. pardus* | Zambia | No | No | 1.5 | 1.3 | M | Yes | | Yes | Yes | Exclude | Exclude | | Patrícia *et al.* 2021 |
| 6352_Zambia | African leopard | *P. p. pardus* | Zambia | No | Yes | 1.0 | 0.2 | M | No | | No | No | Exclude | Exclude | | Patrícia *et al.* 2021 |
| Amurleopard_PPO1 | Amur leopard | *P. p. orientalis* | orientalis | No | No | 33.9 | 27.8 | M | Yes | Yes | Yes | Yes | P_p_orientalis | P_p_orientalis | Yes | Paijmans *et al.* 2021 |
| Amurleopard_PPO5 | Amur leopard | *P. p. orientalis* | orientalis | Yes | No | 33.7 | 27.5 | F | No | Yes | No | No | Exclude | Exclude | | Paijmans *et al.* 2021 |
| SMNH605501 | Amur leopard | *P. p. orientalis* | orientalis | Yes | No | 15.7 | 11.8 | F | No | Yes | No | No | Exclude | Exclude | | Paijmans *et al.* 2021 |
| PP28 | Amur leopard | *P. p. orientalis* | orientalis | No | No | 9.6 | 8.3 | M | Yes | Yes | Yes | Yes | P_p_orientalis | P_p_orientalis | | Paijmans *et al.* 2021 |
| Shinta_10x | Javan leopard | *P. p. melas* | melas | No | No | 17.1 | 11.7 | F | Yes | Yes | Yes | Yes | Exclude | Exclude | Yes | Paijmans *et al.* 2021 |
| MFN_MAM_047501 | Sri Lankan leopard | *P. p. kotiya* | kotiya | No | No | 13.4 | 10.8 | F | Yes | Yes | Yes | Yes | Exclude | Exclude | Yes | Paijmans *et al.* 2021 |
| Bhagya | Indian leopard | *P. p. fusca* | fusca | No | No | 40.3 | 32.4 | M | Yes | Yes | Yes | Yes | Exclude | Exclude | Yes | Paijmans *et al.* 2021 |
| ZMUC29 | Indian leopard | *P. p. fusca* | fusca | No | No | 14.9 | 11.7 | F | Yes | Yes | Yes | Yes | Exclude | Exclude | | Paijmans *et al.* 2021 |
| MFN_MAM_050746 | Indochinese leopard | *P. p. delacouri* | delacouri | No | No | 8.7 | 6.2 | M | Yes | Yes | Yes | Yes | Exclude | Exclude | Yes | Paijmans *et al.* 2021 |

| | | | | | | | | | | | | | | |
|---|---|---|---|---|---|---|---|---|---|---|---|---|---|---|
| MFN_MAM_ 013705 | Indochinese leopard | *P. p. delacouri* | delacouri | No | Yes, transversion only | 5.3 | 3.9 | M | No | Yes | No | Exclude | Exclude | Paijmans *et al.* 2021 |
| Jaguar | Jaguar | *P. onca* | | | | 53.2 | 53.2 | | No | No | No | Exclude | Exclude | Figueiró *et al.* 2017 |
| African_lion | African lion | *P. leo* | | | | 28.1 | 28.1 | | No | No | No | Exclude | Exclude | Cho *et al.* 2013 |
| Snow_leopar d | Snow leopard | *P. uncia* | | | | 28.4 | 28.4 | | No | No | No | Exclude | Exclude | Cho *et al.* 2013 |
| Bengal_tiger | Tiger | *P. tigris* | | | | 28.3 | 28.3 | | No | No | No | Exclude | Exclude | Cho *et al.* 2013 |

**Table S2. Gene ontology and KEGG enrichment result of Zanzibar leopard unique homozygous SNPs**

| #Term | Database | ID | Input number | Background number | P-Value | Corrected P-Value | Input |
|---|---|---|---|---|---|---|---|
| Bile secretion | KEGG PATHWAY | fca04976 | 8 | 68 | 0.000 | 0.037 | AQP4\|ABCG5\|ABCG8\|ADCY9\|AQP8\|ATP1A4\|SLC22A1\|SCTR |
| dynein light intermediate chain binding | Gene Ontology | GO:0051959 | 5 | 14 | 0.000 | 0.031 | CCDC88C\|DNAH3\|DNAH1\|DNAH10\|DNAH5 |
| plasma membrane | Gene Ontology | GO:0005886 | 35 | 754 | 0.000 | 0.031 | AMOTL2\|DUOX1\|SCN9A\|SNAP91\|NOX3\|ATP9B\|SEMA4B\|FYB1\|THSD7B\|ATP6V0A4\|HEPHL1\|ITSN2\|NID2\|NID1\|FCGRT\|GEM\|KCNJ16\|CUBN\|EFNA2\|CRB2\|MCOLN1\|RHOBTB1\|MTNR1B\|ITPR3\|BCAR1\|HOMER2\|KCNT1\|ABCG8\|KCNJ8\|RNPEP\|ITM2C\|SLC19A2\|SIGLEC1\|LRPAP1\|CRHR1 |
| cell-cell adhesion | Gene Ontology | GO:0098609 | 6 | 33 | 0.000 | 0.034 | DCHS2\|CDH23\|FAT2\|FAT1\|DSG2\|DSG4 |
| ATP-dependent microtubule motor activity, minus-end-directed | Gene Ontology | GO:0008569 | 4 | 10 | 0.000 | 0.034 | DNAH3\|DNAH1\|DNAH10\|DNAH5 |
| dynein complex | Gene Ontology | GO:0030286 | 4 | 10 | 0.000 | 0.034 | DNAH3\|DNAH1\|DNAH10\|DNAH5 |
| proteolysis | Gene Ontology | GO:0006508 | 7 | 55 | 0.000 | 0.048 | CASP3\|ERMP1\|CPA5\|CAPN10\|PLAU\|HGFAC\|PRSS38 |
| dynein intermediate chain binding | Gene Ontology | GO:0045505 | 4 | 13 | 0.000 | 0.048 | DNAH3\|DNAH1\|DNAH10\|DNAH5 |

**Table S3. Candidate gene list for body size in canid and coat colour pattern in domestic cats**

| Gene ID | Related phenotype | References |
|---|---|---|
| ADAMTS9 | Canid body size | Plassais et al., 2019 |
| ADAMTSL3 | Canid body size | Plassais et al., 2019 |
| ASCL4 | Canid body size | Plassais et al., 2019 |
| CFA12 (locus) | Canid body size | Plassais et al., 2019 |
| CFA26 (locus) | Canid body size | Hayward et al., 2016 |
| CFA3 (locus) | Canid body size | Plassais et al., 2019 |
| CFA7 (locus) | Canid body size | Plassais et al., 2019 |
| fgf4 | Canid body size | Parker et al., 2009 |
| GHR | Canid body size | Rimbault et al., 2013 |
| HMGA2 | Canid body size | Rimbault et al., 2013 |
| HNF4G | Canid body size | Plassais et al., 2019 |
| IGF1 | Canid body size | Sutter et al., 2007 |
| IGF1R | Canid body size | Hoopes et al., 2012 |
| IGF2BP2 | Canid body size | Plassais et al., 2019 |
| IGSF1 | Canid body size | Plassais et al., 2019 |
| LCORL | Canid body size | Plassais et al., 2019 |
| MITF | Canid body size | Hayward et al., 2016 |
| OGFRL1 | Canid body size | Plassais et al., 2019 |
| R3HDM1 | Canid body size | Plassais et al., 2019 |
| SMAD2 | Canid body size | Rimbault et al., 2013 |
| SMOC2 | Canid body size | Plassais et al., 2019 |
| STC2 | Canid body size | Rimbault et al., 2013 |
| TBX19 | Canid body size | Hayward et al., 2016 |
| THBS2 | Canid body size | Plassais et al., 2019 |
| ZNF608 | Canid body size | Plassais et al., 2019 |
| ASIP | Feline Coat color pattern | Eizirik et al., 2003 |
| MC1R | Feline Coat color pattern | Peterschmitt et al., 2009 |
| DKK4 | Feline Coat color pattern | Kaelin et al., 2021 |
| LVRN | Feline Coat color pattern | Eizirik et al., 2010 |

**Table S4. Body size and coat colour pattern related genes containing homozygous derived alleles in the Zanzibar leopards**

| Chromsome | Position | Reference allele | Alternative allele | Number of African leopard samples (inlcuding the Zanzibar leopard) genotyped | Number of alternative alleles in mainland African leopards | Number of alternative alleles in the Zanzibar leopard | Gene ID | Mutation_type |
|---|---|---|---|---|---|---|---|---|
| chrA1 | 207384247 | G | C | 17 | 5 | 2 | GHR | missense_variant |
| chrA2 | 32104132 | G | A | 18 | 4 | 2 | ADAMTS9 | missense_variant |
| chrA2 | 32119416 | C | T | 17 | 24 | 2 | ADAMTS9 | missense_variant |
| chrA3 | 25149477 | C | A | 17 | 28 | 2 | ASIP | missense_variant |
| chrB1 | 194187148 | C | T | 17 | 26 | 2 | LCORL | missense_variant |
| chrB1 | 194195256 | T | C | 17 | 1 | 2 | LCORL | missense_variant |
| chrB1 | 194195879 | A | G | 17 | 1 | 2 | LCORL | missense_variant |
| chrB1 | 194197712 | C | T | 16 | 19 | 2 | LCORL | missense_variant |
| chrB2 | 153659092 | C | G | 17 | 21 | 2 | SMOC2 | missense_variant |
| chrB2 | 153704388 | C | T | 17 | 2 | 2 | SMOC2 | missense_variant |
| chrB2 | 153704430 | G | A | 17 | 2 | 2 | SMOC2 | missense_variant |
| chrB2 | 154210490 | A | G | 18 | 26 | 2 | THBS2 | missense_variant |
| chrB3 | 3110801 | C | T | 18 | 34 | 2 | ADAMTSL3 | missense_variant |
| chrB3 | 3124941 | T | C | 18 | 19 | 2 | ADAMTSL3 | missense_variant |
| chrB3 | 3164583 | G | C | 18 | 5 | 2 | ADAMTSL3 | missense_variant |
| chrB3 | 3172446 | T | C | 17 | 27 | 2 | ADAMTSL3 | missense_variant |
| chrB4 | 129562193 | A | G | 18 | 33 | 2 | ASCL4 | missense_variant |
| chrC2 | 115196536 | G | A | 16 | 1 | 2 | IGSF10 | missense_variant |
| chrC2 | 115208933 | C | T | 16 | 7 | 2 | IGSF10 | missense_variant |
| chrC2 | 115210145 | C | T | 18 | 1 | 2 | IGSF10 | missense_variant |
| chrC2 | 115210195 | T | A | 18 | 1 | 2 | IGSF10 | missense_variant |
| chrE2 | 63829973 | G | A | 18 | 1 | 2 | MC1R | missense_variant |
